# Gut-immune signaling drives blood-brain barrier damage in pediatric allogeneic stem cell transplant

**DOI:** 10.64898/2026.08.17.745172

**Authors:** Maya R Davies, Courtney B Cross, Feargal J Ryan, Long Yu, Mohsen Dorraki, Zarina Greenberg, Amy Salter, Connor MD Williams, Anna Li, Andrew CW Zannettino, Claudine S Bonder, Cedric Bardy, Hannah R Wardill

## Abstract

Allogeneic hematopoietic stem cell transplantation (allo-HSCT) is a life-saving therapy for children with high-risk hematological diseases. However, allo-HSCT also confers the risk of long-term neurocognitive dysfunction, particularly in pediatric recipients, and the mechanisms underlying this remain poorly understood. While gastrointestinal toxicities and immune responses following allo-HSCT have been well characterized, their contribution to central nervous system toxicities is unknown. Here, using clinical biomarker analysis, we show evidence of blood-brain barrier (BBB) dysfunction in pediatric allo-HSCT, associated with IL-6 signaling and reduced levels of brain-derived neurotrophic factor. Pre-transplant gastrointestinal mucosal barrier injury was associated with post-transplant BBB leakage, implicating disrupted gut-brain-axis signaling. *In vitro*, gut damage-associated immune activation induced apoptosis and remodeling of brain microvascular endothelial cells (BMECs), with surviving cells exhibiting tight junction disruption and cytoskeletal reorganization. Plasma from allo-HSCT recipients similarly induced BMEC apoptosis. Notably, both immune signaling- and patient plasma-induced BMEC apoptosis were prevented by IL-6 inhibition or supplementation with the gut microbiota-derived metabolite propionate. Together, these findings identify immune signaling as a correlate of BBB damage clinically and a causative driver *in vitro* in pediatric allo-HSCT.

## Introduction

Hematopoietic stem cell transplant (HSCT) has revolutionized the outcomes of treatment-resistant and refractory hematological malignancies. These diagnoses were once terminal but with advances in HSCT, they now claim a 70% 5-year survival rate – an increase of 40% over the last five decades (1, 2). Despite these improvements in ***overall survival***, HSCT is associated with significantly reduced ***quality of life*** which is attributable to the aggressive nature of this therapy (3–5). HSCT involves high-dose myeloablative conditioning (chemotherapy ± total body irradiation) to eradicate tumor cells and ablate the host’s own immune system in preparation for subsequent stem cell grafting (6). Allogeneic HSCT (allo-HSCT) is often preferred, as donor-derived T cells can mount a graft vs tumor (GvT) response to enhance anti-cancer efficacy while reconstituting immune function (6). However, in allo-HSCT, clinical benefit demands a delicate balance between efficacy and safety, with treatment-related mortality in some studies exceeding that from underlying disease progression (7–9). Myeloablative conditioning-related infection and alloreactive immune responses against healthy recipient tissues (termed graft vs host disease, GvHD), remain leading causes of non-relapse mortality in this patient population (7–9).

HSCT survivors, particularly children, are also highly vulnerable to long-term sequelae, with up to 90% developing treatment-related complications (10, 11). These effects span multiple organ systems, including gastrointestinal, skeletal, pulmonary, hepatic, renal, and central nervous system (CNS) involvement (12). Neurocognitive impairment is frequently reported among allo-HSCT survivors (11-59% prevalence), particularly for pediatric recipients, affecting domains of attention, memory, executive function, processing speeds, often cumulating in reduced IQ (13–16). Post-mortem analysis identified neuropathological abnormalities in up to 91% of HSCT survivors, underscoring substantial neurotoxic effects (17). Despite this burden, the mechanisms underlying these neurocognitive sequelae remain largely unexplored and hence, methods to prevent or manage these cognitive side effects are lacking.

When considering systemic influences of CNS function, the importance of the blood-brain barrier (BBB) cannot be overlooked. The blood-brain barrier (BBB) is an intricate, multi-cellular structure which serves as a crucial interface between systemic circulation and the highly vulnerable brain parenchyma, playing a critical role in maintaining CNS homeostasis (18). Disruption or dysfunction of the BBB is therefore a key pathological feature shared by numerous neurological conditions, including multiple sclerosis, Parkinson’s disease, Alzheimer’s disease, and COVID-19 brain-fog (19, 20). As brain function is inseparable from the function of the cerebrovascular system, interventions aimed at modulating or reinforcing the BBB are increasingly recognized as promising neuroprotective strategies (19, 21). However, investigation of the impact of allo-HSCT on BBB function has not previously been performed.

In recent years, research which leverages on the complex interactions between the immune system and the resident microbes of the gastrointestinal system – the gut microbiota – has shown significant promise to modulate immune responses in allo-HSCT and improve outcomes in this population (22). In a similar manner, the immune system and the gut microbiota have emerged as key interacting mediators of BBB integrity and CNS function, via exerting inflammatory/damaging or immunosuppressive/protective effects (23–25). Accordingly, targeting microbiota-immune interactions serves as a powerful strategy to regulate BBB permeability and limit CNS exposures to noxious circulating factors (20, 26–30). However, in allo-HSCT, the impact of microbiota-immune signaling on the CNS has not previously been investigated. Here, we present a bed-to-bench investigation of BBB function in pediatric allo-HSCT, providing the first clinical evidence of barrier leakage coupled with *in vitro* characterization of immune-mediated endothelial apoptosis and remodeling. We also demonstrate that targeting pro-inflammatory signaling and microbial metabolism is effective in preventing direct allo-HSCT plasma-induced apoptosis of brain microvascular endothelial cells, suggesting there is therapeutic potential of leveraging microbiota-immune interactions.

## Results

Plasma samples from N=47 pediatric allo-HSCT recipients (<18-years-of-age at time of transplant, Table 1) were sourced from the CRYOSTEM biobank (France; project no. CS-23-02). Median age at time of HSCT was 11.66 years and with a majority of male patients (66%). The predominant indication for HSCT was leukemia, acute myeloid (28%) or acute lymphoid (21%). Conditioning treatments for HSCT involved various myeloablative techniques: 30% of patients received total body irradiation and common chemotherapies administered (as part of combination therapies) included busulfan (53%), cyclophosphamide (49%) and fludarabine (40%). HSCT-related complications were the predominant cause of mortality in the cohort (63.6%, 7/11), with reports of relapse/progression-related mortality in only 4 patients (8.5%, 4/47).

**Table 1.**
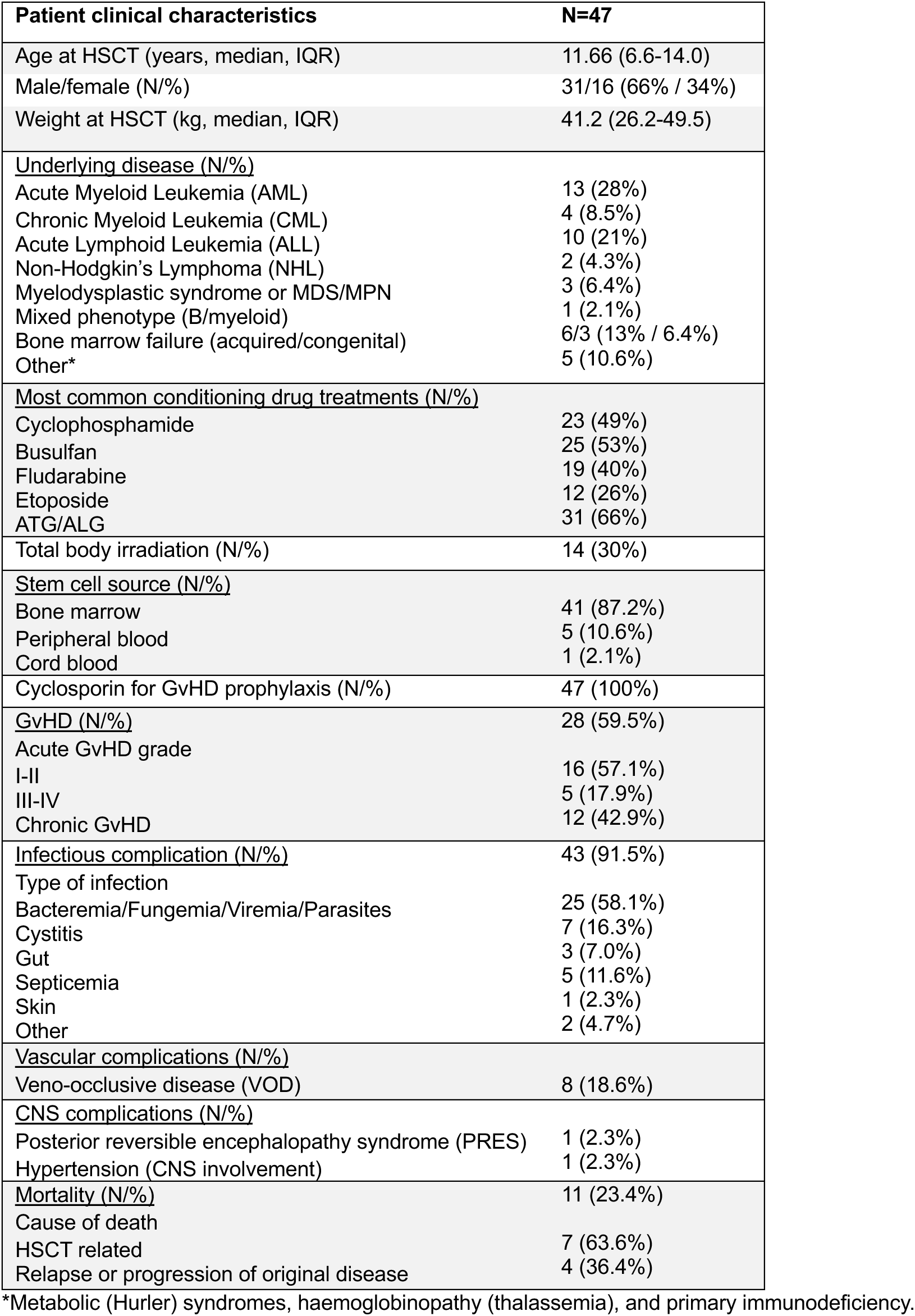
Clinical characteristics of allo-HSCT recipients.

### Peripheral immune activation and disruption of the gut microenvironment in allo-HSCT

A baseline, pre-HSCT (and pre-conditioning) sample and a post-HSCT sample (time varied, median: 98.5-days post-HSCT) were sourced from each patient, with additional follow-up samples for some patients (Fig.1A). To assess the general effects of allo-HSCT, pre- and post-HSCT samples were compared. Quantification of circulating peripheral cytokines revealed significant increases in several cytokines (Fig.1B); IL-6 demonstrated the largest relative increase from baseline (2.6-fold, P<0.0001) among other upregulated pro-inflammatory mediators: IL-5 (1.8-fold, P<0.0001), TNFα (1.8-fold, P=0.0003), and IFNγ (1.3-fold, P=0.0006).

**Fig. 1.**
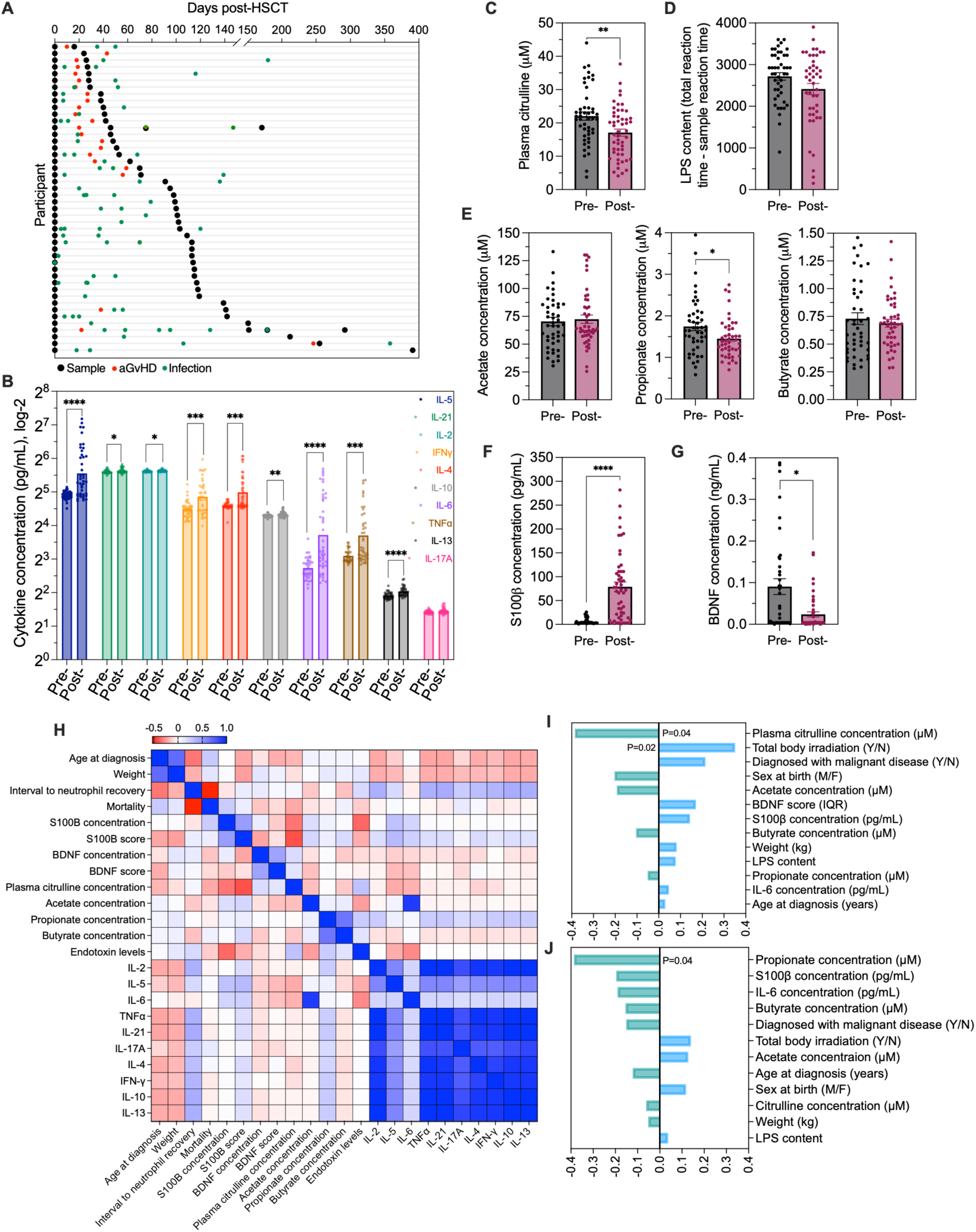
Allo-HSCT biomarker characterisation and correlation to poor neurological outcomes. (A) N=47 pediatric HSCT recipients with baseline (pre-HSCT) and post-HSCT sample (black circles). Time of acute GvHD (aGvHD) diagnosis is indicated with a red circle, and green circles indicate onset of an infectious event (Supplementary Fig.1: GvHD sub-group analysis and longitudinal analysis for figures B-G). (B-G) Quantification of plasma levels of key biomarkers. (B) Cytokines pre- and post-HSCT determined via multi-analyte array (multiple Mann-Whitney tests). (C) Citrulline analyzed via mass spectrometry (unpaired t-test). (D) Endotoxin (lipopolysaccharide, LPS) content measured via colorimetric assay. (E) Mass spectrometry quantification of short chain fatty acids (SCFAs): acetate, propionate, butyrate. (F-G) ELISA results of (F) S100β and (G) BDNF concentrations (measured via ELISA; Mann-Whitney test for D-G). (H) Correlation matrix (displayed in heatmap form for R values) indicating strength and direction of linear and temporally matched relationships between biological variables and clinical factors (Supplementary Fig. 2 for Pearson’s correlation analysis for statistical significance). Given the exploratory nature of the analysis and the study sample size, P-values were interpreted descriptively without formal adjustment for multiple comparisons. (I-J) Baseline (pre-HSCT) factors associated with (I) post-HSCT BBB leakage (4^th^ IQR S100β concentrations), or (J) post-HSCT reductions in BDNF (1^st^ IQR BDNF concentrations). Data depicts the R-value for each factor (I-J Pearson’s correlation). *p<0.05, **p<0.01, ***p<0.001, ****p<0.0001. Histogram data represent mean ±SEM. Matrix color demonstrates strength and direction of data. Bar charts are a visualization of the correlation co-efficient for each variable.

Consistent with gastrointestinal injury, pediatric allo-HSCT recipients demonstrated significantly reduced plasma citrulline concentrations following transplant (1.8-fold, P=0.003; Fig.1C). Citrulline is a clinically validated biomarker of mucosal (gastrointestinal) barrier injury, a common occurrence during HSCT as a consequence of conditioning-induced mucositis and disruption of the gut microbiota (31). Mucosal barrier injury is closely intertwined with immune activation, as the gut constitutes the body’s largest immunological organ and serves as a critical barrier separating luminal microbes from systemic circulation (32). While circulating levels of LPS, a bacterial endotoxin broadly associated with gut microbial disruption, were not significantly altered (P=0.12; Fig.1D), analysis of circulating short-chain fatty acids (SCFA) revealed significantly reduced concentrations of the neuroprotective metabolite propionate (1.2-fold; P=0.02, Fig.1E).

### Increased BBB permeability following pediatric allo-HSCT

To assess BBB function clinically, we quantified plasma concentrations of S100β, an astrocyte-derived protein too large (21kDa) to extravasate the intact BBB and hence is used as a biomarker of BBB disruption – S100β predicts computed tomography (CT) results in mild traumatic brain injury and is incorporated into Scandinavian clinical guidelines to reduce unnecessary CT usage (33, 34). Post-allo-HSCT, circulating S100β levels increased 10-fold from baseline (pre=5.7pg/mL; post=78.8pg/mL; P<0.0001; Fig.1F), consistent with increased BBB permeability. As a complementary neurological marker, we also measured plasma levels of brain-derived neurotrophic factor (BDNF), a predictor of cognitive function in cancer survivor cohorts (35–38). BDNF was significantly reduced in the post-HSCT samples (a 3.2-fold decrease; P=0.04; Fig.1G) which was most profound within the first 50-days-post-transplant (P=0.01; Supplementary Fig.1O). We next performed exploratory analysis of relationships between biological factors and clinical variables via Pearson correlation matrix analysis (Fig.1H and Supplementary Fig. 2A). We observed (1) S100β (both concentration and interquartile range [score]) negatively correlated with citrulline (R=-0.34; P<0.01 and r=-0.42; P<0.001, respectively), and (2) S100β score positively correlated with circulating IL-6 concentrations (R=0.23; P<0.05). In addition, S100β score was the sole factor which demonstrated a significant (negative) association with BDNF concentrations (R=-0.23; P<0.05), supporting the importance of BBB integrity for CNS function (39–42). Elevated S100β concentrations were significantly correlated with increased mortality (R=0.25; P<0.05), and high S100β levels post-allo-HSCT predicted reduced overall survival on Kaplan-Meier curves (P<0.05; Supplementary Fig. 2B) and while univariable Cox regression analysis suggested higher risk of death (HR=2.4, 95% CI 0.73-9.34), this was not statistically significant.

### Pre-HSCT gut microenvironmental factors are associated with post-transplant neurovascular biomarkers

We next performed exploratory feature prioritization between baseline (pre-HSCT) factors and post-transplant outcomes of interest: (1) BBB damage (high plasma S100β) and (2) low BDNF, using Pearson’s correlation analysis. Reduced plasma citrulline prior to allo-HSCT (indicative of mucosal barrier injury) emerged as the strongest factor at baseline associated with post-transplant BBB damage, as defined by S100β concentrations within the highest inter-quartile range (IQR, R= −0.4; P=0.02; Fig. 1I). Inclusion of total body irradiation in the conditioning therapy was also significantly associated with subsequent increases in S100β (R=0.35; P=0.04), which aligns with the previously published barrier disruptive consequences of irradiating therapies (43–45). Reduced propionate levels was the sole baseline factor associated with post-HSCT BDNF depletion (R= −0.4; P=0.04; Fig.1J). Together, these exploratory findings implicate pre-transplant gastrointestinal mucosal integrity and microbial metabolism in subsequent neurovascular dysfunction following allo-HSCT.

### HSCT conditioning, and microbial- and inflammatory-insults are directly toxic to iPSC-derived BMECs

To investigate toxicities mechanistically we next sought to replicate conditioning chemotherapy-induced mucosal barrier injury *in vitro*. We performed a dose-finding experiment – exposing human colonic epithelial T84 cells to the active metabolite of cyclophosphamide (phosphoramide mustard, PM), a commonly used conditioning chemotherapy, for 48-hours. 1.28mM PM was required to induce >10% loss of viability (P<0.0001; Supplementary Fig.3A, 3B), a dose which we confirmed induced mucosal barrier injury via caspase-3 immunofluorescent staining of T84 monolayers on transwells (P<0.01; Supplementary Fig.3C, 3D). We next examined the effects of dual treatment of PM and LPS on human primary peripheral blood mononuclear cells (PBMCs, Supplementary Fig.3E). A 24-hour exposure to 1.28mM PM and 10ng/mL LPS resulted in a significant reduction in PBMC viability (P<0.01; Supplementary Fig.3F) and robust induction of IL-6 secretion, whereas IL-6 was undetectable in PBS-treated control supernatant (Supplementary Fig.3G).

We next looked to characterize effects on iPSC-derived BMECs monolayers cultured in transwells. To initially capture the relevance of this model, iPSC-derived BMECs were compared with primary human BMECs, human umbilical vein endothelial cells (HUVECs), and T84 colonocytes at transcriptomic, structural, and functional levels (Supplementary Fig.4). We performed bulk RNA sequencing on these cells (treatment naïve); iPSC-derived BMECs from different sources clustered together and, while remaining transcriptomically distinct from primary BMECs (as reported previously) (46), demonstrated superior barrier functionality and tight junction organization compared with HUVEC comparators. We then investigated direct effects of PM on iPSC-derived BMECs and also of LPS, given the established barrier-damaging effects of the bacterial endotoxin (Fig.2A). Exposure to 1.28mM PM for 48-hours significantly increased barrier permeability (112%), quantified by greater translocation of 4kDa FITC across the BMEC monolayer (P=0.002; Fig.2B). PM also exerted directly cytotoxic effects on BMECs, with 1.7x greater caspase-3+ area (P=0.002; Fig.2C, 2D), and induced morphological changes, with increased endothelial cell size (2.5-fold, P=0.0005; Fig.2E) and cytoskeletal F-actin reorganization (Fig.2C). Both LPS and PM disrupted BMEC tight junctions, distinguished by disorganized ZO-1 staining with poor membrane specificity (P<0.05; Fig.2F, 2G). Importantly, prior *in vivo* research demonstrates ZO-1 inhibition alone is sufficient to enable access of chemotherapeutic agents (doxorubicin and paclitaxel) to the CNS, in the context of delivering anti-cancer therapy to the brain (47). Consistent with this, our findings demonstrate that cyclophosphamide-based conditioning regimens disrupt ZO-1 junctions and cause substantial endothelial toxicity, which may result in increased off-target exposure of the CNS to cytotoxics.

**Fig. 2.**
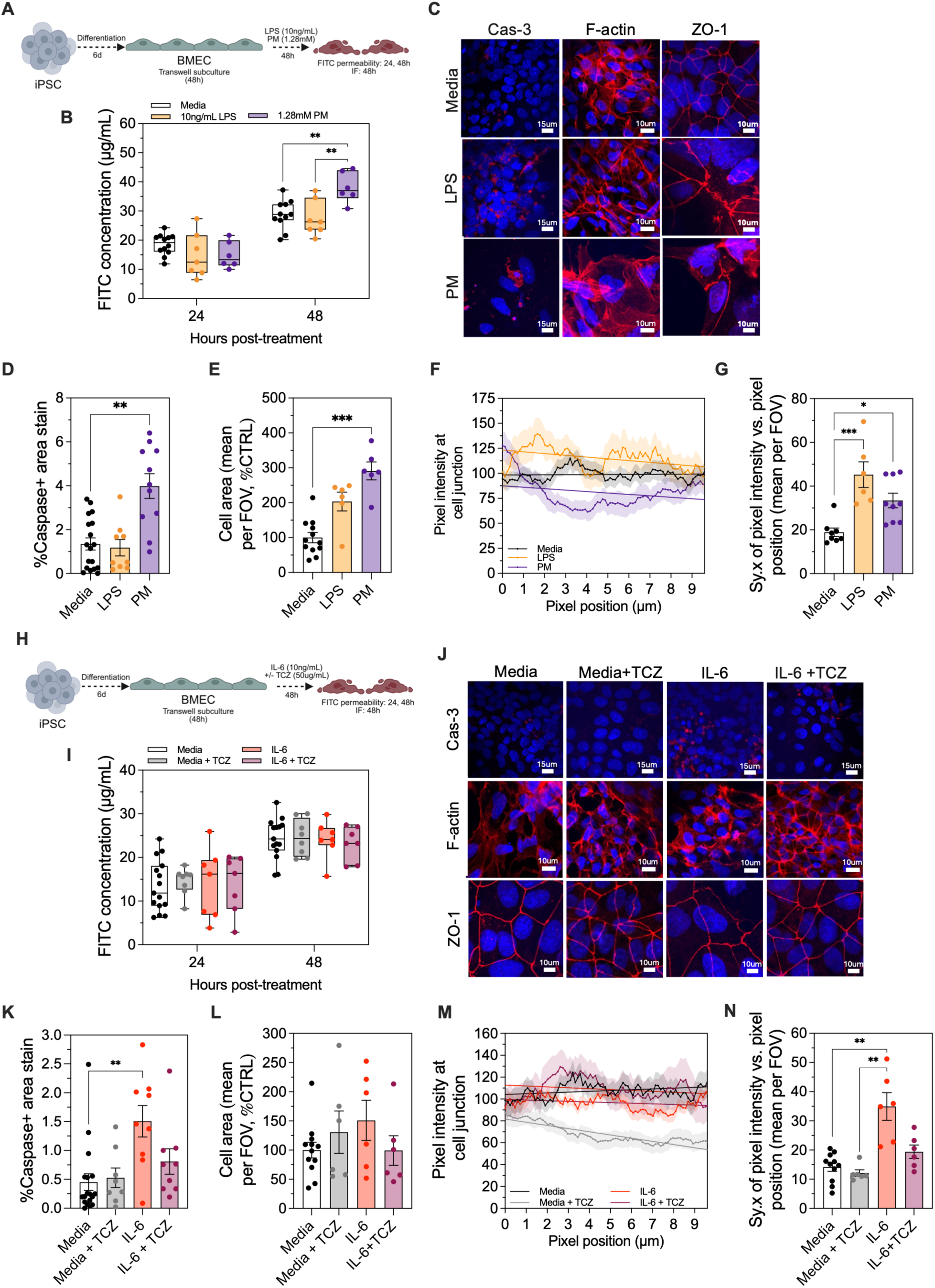
Active metabolite of cyclophosphamide, bacterial endotoxin, and IL-6 disrupt iPSC-derived BMEC monolayers. (A) Experimental design of 48-hour exposure of iPSC-derived BMECs to 1.28mM PM, 10ng/mL LPS, or normal media control while cultured as monolayers in transwells. Functional assays were performed on all transwells across two independent experiments, which were then assigned to different staining conditions resulting in N≥6 transwells for FITC analysis (B,I) and N≥6 fields of view (FOVs) across two transwells for immunofluorescence analysis (D-G, K-N). (B) Quantification of 4kDa FITC translocation across BMEC monolayer, from apical to basolateral transwell compartment, at 24- and 48-hours following treatment initiation (mixed-effects analysis). (C) Representative images of immunofluorescent analysis of caspase-3 (cas-3), F-actin, and ZO-1 48h post-treatment. (D) Quantification of caspase-3 %area positive staining per FOV. (E) Measurement of mean cell standard area per field of view (FOV; expressed as % of control). (F) Consistency of ZO-1 staining intensity along 10μm of the linear junction, with the x-axis representing a position along the junction and the y-axis indicating the pixel fluorescent intensity at the position, line of best fit plotted by linear regression (combined results for all images per condition) (G) Quantification of xy plots in (F) for individual FOVs as standard deviation of individual points from the regression line (sy.x), expressed as mean per FOV. (Kruskal-Wallis test for D,E,G). (H) Experimental design of 48h exposure of iPSC-derived BMECs to 10ng/mL IL-6, 10ng/mL IL-6 + 50μg/mL tocilizumab (TCZ), media +50μg/mL TCZ, and media only control. (I) Quantification of 4kDa FITC translocation across BMEC monolayer, from apical to basolateral transwell compartment, at 24 and 48h following treatment with IL-6 (±TCZ) (mixed-effects analysis). (J) Representative images of caspase-3, F-actin, and ZO-1 immunofluorescent analysis at 48h. (K) Caspase-3+ %area stain. (L) Mean cell standard area per FOV. (M) Consistency of ZO-1 staining intensity along 10μm of the linear junction, with line plotted via linear regression (all images per condition combined for representative figure). (N) Quantification of (M) for each FOV individually as standard deviation of individual points from the regression line (sy.x), expressed as mean per FOV. (Kruskal-Wallis test for K,L,N). *p<0.05, **p<0.01, ***p<0.001. Histogram data represent mean ±SEM. Box and whisker plots represent median and min-max values, xy charts demonstrate mean ±SEM.

We next tested the effects of IL-6 signaling on BMEC monolayers as this cytokine was profoundly elevated in the patient plasma post-allo-HSCT. To confirm direct IL-6 effects, we co-administered the clinically approved IL-6-receptor monoclonal antibody tocilizumab (TCZ; Fig.2H). While IL-6 did not significantly alter 4kDa FITC permeability across BMEC monolayers (Fig.2I), it did induce significant endothelial toxicity as evidenced by increased caspase-3 induction (3x greater positively stained area, P=0.003; Fig.2J, 2K). IL-6 did not produce overt cytoskeletal reorganization or endothelial swelling (Fig.2J, 2L) but caused significant disruption of ZO-1 tight junction architecture (P<0.0001; Fig.2M, 2N), consistent with previous studies (48, 49). Importantly, TCZ attenuated both apoptosis and tight junction disruption, confirming IL-6-dependent endothelial injury.

### Peripheral immune activation drives BMEC injury and remodeling

Having established the direct effects of conditioning-, microbial- and inflammatory insults, we next modelled their integrated effects through peripheral immune activation. Specifically, conditioned medium (CM) from PBMCs stimulated with PM and LPS (stim-PBMC-CM), and consequently enriched for IL-6, was diluted 1:2 in normal BMEC media and applied to the apical compartment of BMEC cultures for 48-hours (Fig.3A). This was compared to the effects of CM from PBMCs treated only with PBS (ctrl-PBMC-CM) and also a media only control. Exposure to the combined insults of the stim-PBMC-CM recapitulated and amplified the effects observed in experiments using isolated factors, with formation of F-actin stress fibers (Fig.3B), increased caspase-3+ staining (2.9-fold, P=0.0006; Fig.3C), cell swelling/oedema (4.5x increased cell size, P<0.0001; Fig.3D), and extensive ZO-1 tight junction disorganization (P=0.01; Fig.3E, 3F). No significant alterations were observed following ctrl-PBMC-CM exposure, indicating that BMEC injury was driven by factors released following PBMC activation.

**Fig. 3.**
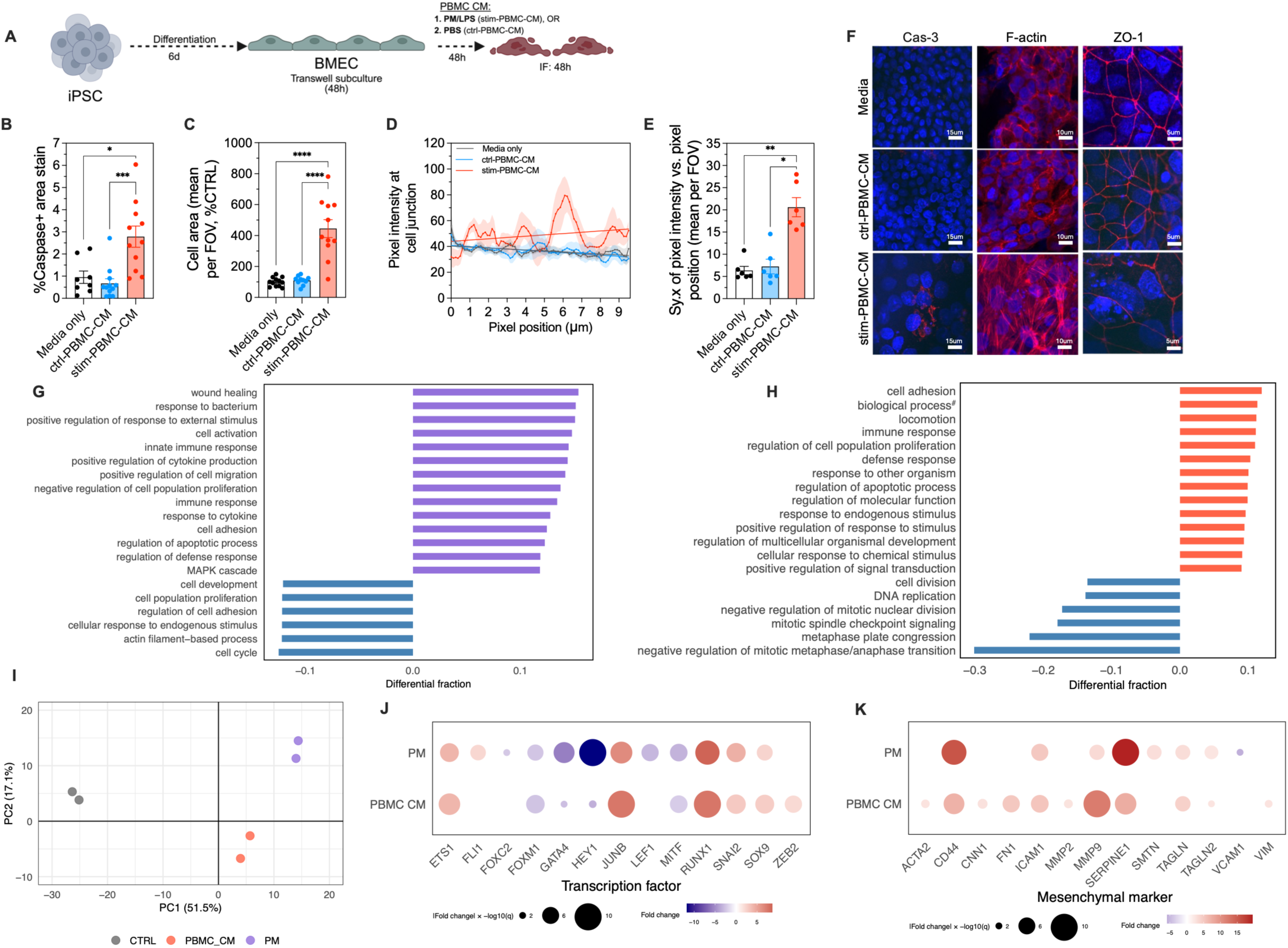
Peripherally derived inflammatory signaling induces significant BMEC stress and remodeling. (A) Experimental design of 48-hour exposure of iPSC-derived BMECs to 50% conditioned media (CM) from 1.28mM PM + 10ng/mL LPS stimulated (stim-PBMC-CM), or vehicle control-treated PBMCs (ctrl-PBMC-CM), or normal media control for 48-hours (N≥8 FOVs across 4 transwells over two independent experiments for each immunofluorescent stain). (B) Representative images of immunofluorescent staining for caspase-3, F-actin, and ZO-1 after 48-hours. (C-F) Quantification (C) caspase-3-positive area, (D) mean cell area per FOV, and ZO-1 junctional continuity assessed by linear regression of staining intensity along 10μm junctional segments (E), with variability expressed as the standard deviation from the regression line (sy.x) per FOV (F) (Kruskal-Wallis test). (G-H) Gene enrichment analysis for significant up and downregulated pathways using gprofiler2 summarized under parent terms using rrvgo for BMECs exposed to (J) 1.28mM PM, or (K) stim-PBMC-CM (^#^involved in interspecies interaction between organisms) relative to media only control (N=5 transwells pooled into N=2 replicates for bulk RNA sequencing). (I) Principal component analysis for media (CTRL), PM, and stim-PBMC-CM. (J) Dotplot for expression of key transcription factors implicated in EndoMT for PM and stim-PBMC-CM, where dot color represents fold-change (relative to control) and dot size represents: fold_change × −log10(q-value) to avoid infinite values. (K) Dotplot for expression of key mesenchymal genes implicated in EndoMT for PM and stim-PBMC-CM, where dot color represents fold-change (relative to control) and dot size represents: fold_change × −log10(q-value) to avoid infinite values. *p<0.05, **p<0.01, ***p<0.001. Histogram data and the xy chart represent mean ±SEM.

With clear phenotypic disruption observed, we next performed bulk RNA sequencing on BMECs exposed to (1) stim-PBMC-CM, (2) BMECs exposed to PM, and (3) media-only control BMECs to confirm an altered cellular identity. For each condition, RNA from N=5 transwells were pooled into N=2 sequencing replicates, each generating ~47 million paired-end reads. Pathway enrichment analysis with gprofiler2 and parent term summation via the rrvgo package identified enriched pathways for immune response/inflammation, apoptosis, cell activation and defense response following PM exposure (Fig.3G). Concurrently, pathways associated with cell function, development, adhesion and actin processes were depleted. In BMECs exposed to stim-PBMC-CM, enriched pathways pertained to cell adhesion, immune response, defense response, apoptosis, and cellular signaling, while pathways involved in cell division were depleted (Fig.3H). Principal component analysis revealed clear separation between treatment groups (Fig.3I). Consistent with endothelial phenotypic transformation, genes associated with endothelial-to-mesenchymal transition (EndoMT), a canonical endothelial transdifferentiation pathway, were broadly upregulated, including EndoMT transcription factors (50) and key mesenchymal markers (50, 51) in particular with stim-PBMC-CM exposure (Fig.3J, 3K). Together, these phenotypic and transcriptional changes demonstrate profound endothelial stress and remodeling, consistent with partial loss of endothelial identity.

### IL-6 inhibition and propionate supplementation prevent immune-mediated BMEC apoptosis, but not endothelial remodeling

To investigate the direct role of IL-6 in the BMEC pathology induced by peripheral immune activation, BMEC monolayers were exposed to stim-PBMC-CM ± IL-6-receptor antibody, tocilizumab (TCZ; Fig.4A). Exposure to the inflammatory stim-PBMC-CM resulted in a modest but statistically significant reduction in 4kDa FITC-dextran translocation across the BMEC monolayer (−1.28-fold, P=0.04; Fig.4B). However, this was likely confounded by pronounced endothelial cell swelling (3.4-fold increased cell size, P<0.05; Fig.4E) which could reduce paracellular flux by reducing paracellular zones rather than reflecting improved barrier function, particularly as stim-PBMC-CM also induced the consistent phenotype of caspase-3 activation (~13-fold increased stain area, P=0.0001; Fig.4D). Importantly, co-administration of TCZ effectively prevented this profound stim-PBMC-CM-induced BMEC apoptosis, implicating IL-6 signaling as a key mediator. In contrast, TCZ did not mitigate the other cellular pathological features: endothelial cell swelling and F-actin cytoskeletal reorganization persisted despite IL-6 inhibition, suggesting these phenotypes are regulated by IL-6 independent pathways. Moreover, although IL-6 alone induced ZO-1 endothelial tight junction disruption which was inhibited by TCZ (Fig.2N), TCZ was insufficient to protect against tight junction disruption induced by the complex inflammatory milieu of the conditioned media (Fig.4F, 4G).

**Fig. 4.**
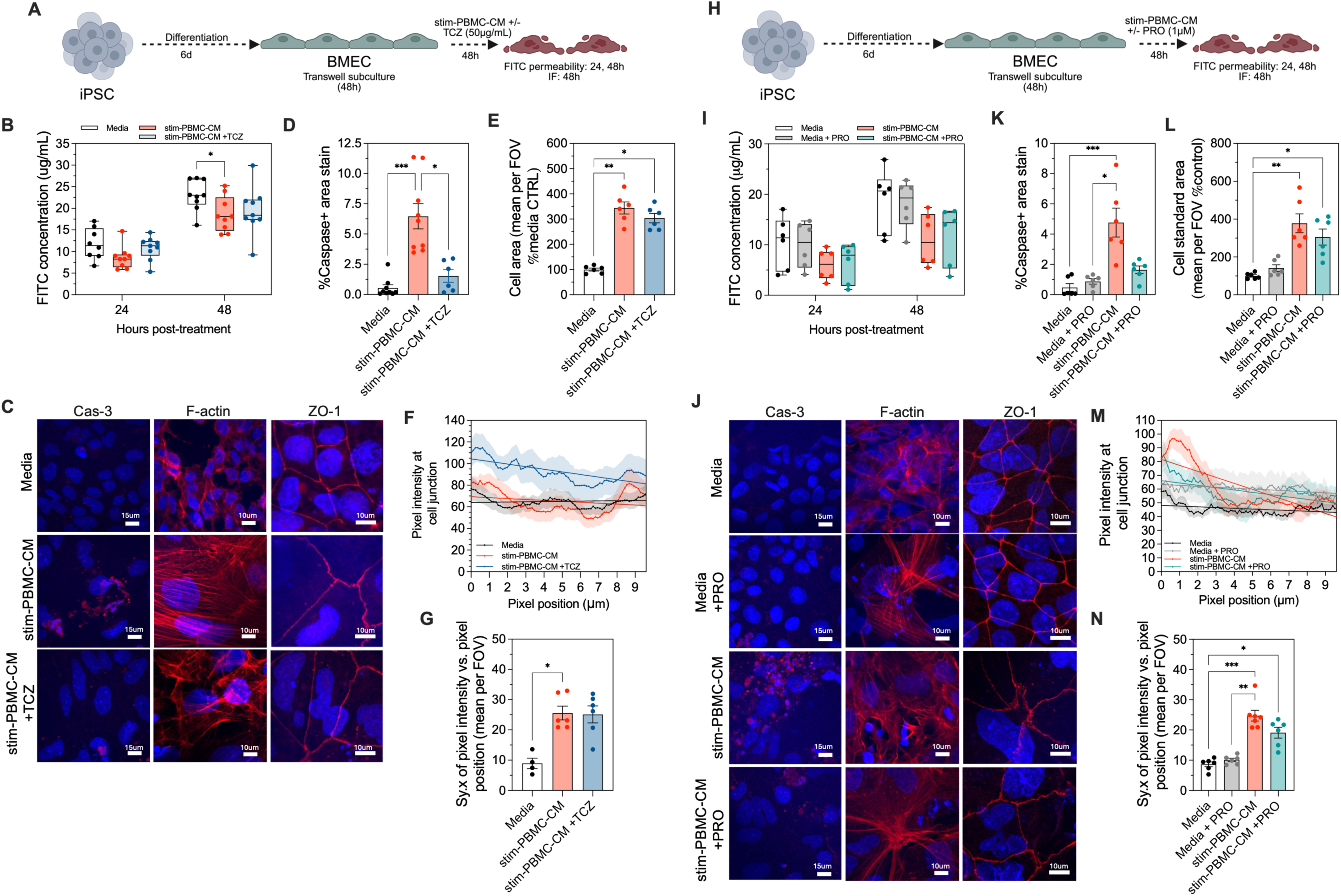
Anti-inflammatory interventions prevent BMEC apoptosis but not tight junction disruption. (A) BMEC exposure to stim-PBMC-CM (50% dilution) for 48h ±TCZ (50μg/mL), or media-only control. N≥6 transwells across two independent experiments for FITC analysis (B, I) then assigned to different stains for N≥6 FOVs across two transwells for immunofluorescence analysis (D-G, K-N). (B) 4kDa FITC-dextran permeability measured at 24 and 48h post-treatment initiation (mixed-effects analysis). (C) Representative immunofluorescence images of caspase-3 (Cas-3), F-actin, and ZO-1 at 48h. (D-G) Quantification of (D) caspase-3 positive area, (E) mean cell area per FOV, and ZO-1 junctional continuity assessed by linear regression of staining intensity over 10μm junctional segments (F), with variability expressed as the standard deviation from the regression line (sy.x) per FOV (G) (Kruskal-Wallis test). (H) Experimental schematic of BMEC exposure to stim-PBMC-CM (50%) ±PRO (1μM), or media-only ± PRO controls, for 48h. (I) FITC-dextran permeability 24 and 48h following PRO treatment (mixed-effects analysis). (J) Representative immunofluorescence images of Cas-3, F-actin, and ZO-1 at 48h. (K-N) Quantification of (K) caspase-3 positive area, (L) mean cell area per FOV, and (M,N) ZO-1 junction formation continuity as described in (F,G) (Kruskal-Wallis test). *p<0.05, **p<0.01, ***p<0.001. Histogram data and xy charts represent mean ±SEM. Box and whisker plots represent median and min-max values.

Given the targeted mechanism of TCZ, limited to isolated blockade of IL-6 signaling, we next explored the effects of propionate supplementation (Fig.4H) – a broadly anti-inflammatory microbial metabolite predictive of post-transplant BDNF levels in our biomarker analysis. Again, the stim-PBMC-CM exposure appeared to reduce flux of FITC-dextran across the BMEC monolayer (although this did not reach statistical significance; Fig.4I), once more accompanied by endothelial swelling (278% increased cell area, P<0.01; Fig.4J, 4L). Propionate effectively prevented the CM-induced increase in caspase-3 (2.9-fold reduction of %area stain than with stim-PBMC-CM alone and P=ns vs. media control; Fig.4K), supporting a role for inflammation-driven apoptotic signaling. However, propionate did not rescue the other pathological features of F-actin stress fiber formation (Fig.4J), increased cell area (Fig.4I), and ZO-1 tight junction disorganization (Fig.4M, 4N).

### Allo-HSCT plasma directly induces BMEC toxicity *in vitro*

Next we applied plasma from pediatric allogeneic HSCT recipients directly to our *in vitro* model for 24-hours (Fig.5A). Plasma from 10 patients with the most extreme concentrations of citrulline were pooled into two groups (high or low, N=5/group) and diluted 1:4 in BMEC medium (with the addition of penicillin-streptomycin antibiotics to minimize the risk of contamination). As no significant differences were observed between the low- and high-citrulline groups (Supplementary Fig.5), all allo-HSCT plasma-treated wells were combined for overall analyses. Patient-derived allo-HSCT plasma significantly increased levels of BMEC apoptosis (18x increased %caspase-3+ area) relative to media (media+Ab) control (P<0.05; Fig.5B, 5C). In contrast, plasma from a healthy control did not induce significant apoptosis, supporting the specificity of the effect to the allo-HSCT circulating milieu. Parallel administration of either propionate or TCZ prevented allo-HSCT plasma-induced BMEC apoptosis, with %caspase-3+ staining comparable to healthy plasma or media controls.

**Fig. 5.**
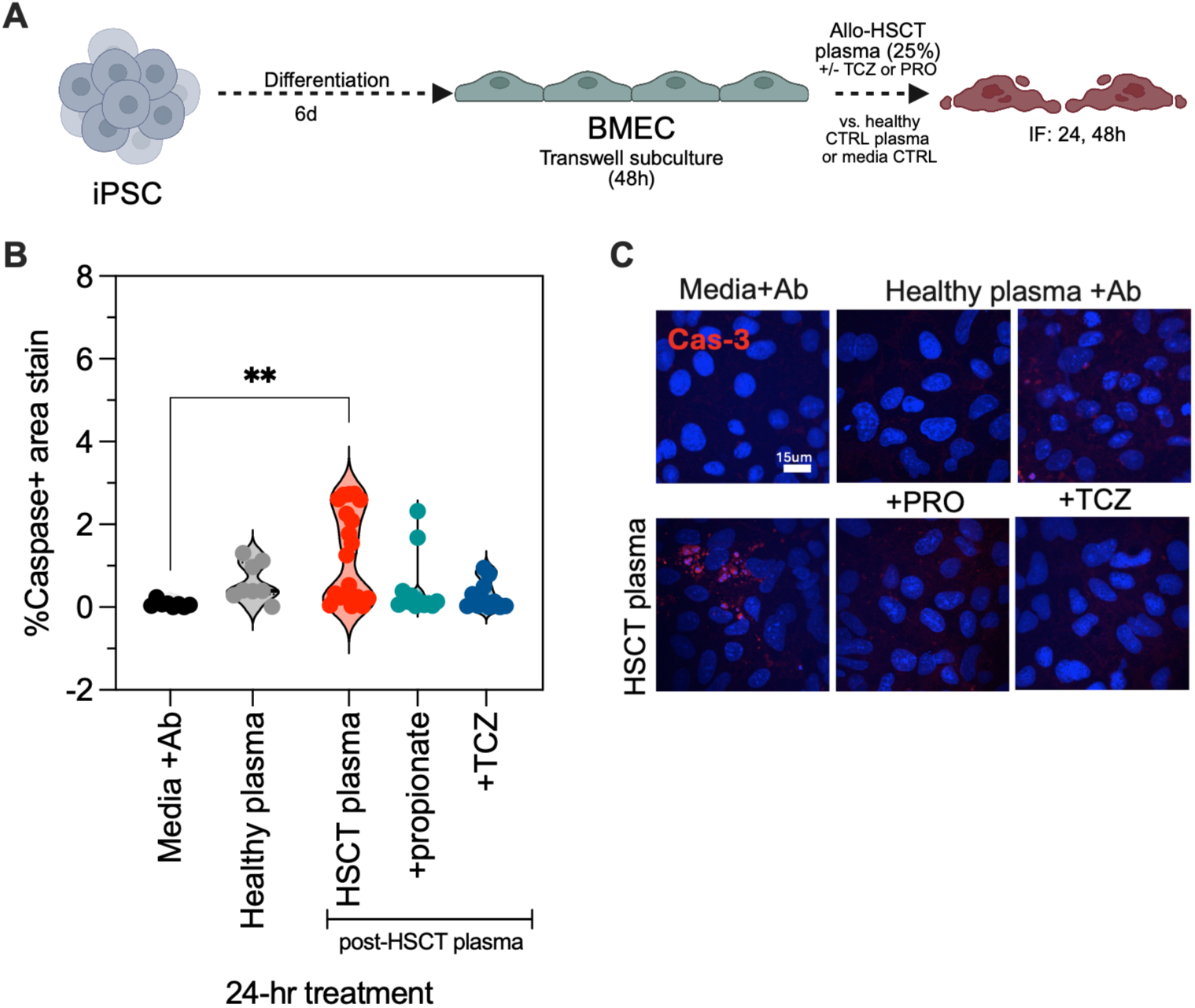
Allo-HSCT patient plasma induced BMEC cell toxicity, prevented by propionate or TCZ. (A) Experimental design of exposing BMEC monolayers to allo-HSCT patient plasma. Plasma was pooled from N=10 patients (post-transplant time-point) and diluted 1:4 in normal endothelial media ± interventions (50μg/mL TCZ or 1μM propionate). This was compared to plasma from a healthy control or media only control. Antibiotics (Ab) was added to all conditions. N=2 transwells for media+Ab and healthy plasma and N=4 allo-HSCT plasma, allo-HSCT plasma + TCZ, and allo-HSCT + propionate (across two independent experiments). (B) Caspase-3 positive area determined via immunofluorescent analysis (Kruskal-Wallis test). (C) Representative images quantified in (B). **P<0.01. Violin plots demonstrate median and min-max values.

## Discussion

Despite its clinical significance, neurocognitive impairment caused by cancer therapies remains poorly understood, profoundly impacting the quality of cancer survival. Whilst involvement of the BBB has been theorized, there has been limited investigation of underlying mechanisms. Our findings provide clinical evidence consistent with BBB dysfunction following pediatric allo-HSCT, evident within the first 100 days post-transplant. To our knowledge, this represents the first clinical evidence of BBB dysfunction in this setting, with prior evidence limited to preclinical studies demonstrating the presence of allogeneic T cells within the CNS following HSCT (52). Given limited prior mechanistic investigation, a particularly novel aspect of our work is the in-depth characterization of BMEC toxicity. A key finding was the consistent emergence of a clear morphological signature of endothelial stress in response to cytotoxic and pro-inflammatory insults, with increased BMEC size, cytoskeletal remodeling, and tight junction disruption. Importantly, this represents the first demonstration of a causal link between peripheral inflammation and cerebral vascular injury in the context of allo-HSCT.

Endothelial cells are inherently plastic in nature and can undergo extensive cellular remodeling in response to injury, either as an adaptive response or a maladaptive survival strategy which compromises cellular function (51). Consistent with this paradigm, BMEC exposure to conditioning, and microbial- and immune-associated insults induced F-actin stress fiber formation and/or ZO-1 disorganization, accompanied by transcriptional changes consistent with cellular stress and partial loss of endothelial identity. Similar BMEC morphological changes have been reported in the context of Alzheimer’s disease-associated BBB damage: cytokine stimulation of BMECs generated from patient-derived iPSCs induced cytoskeletal remodeling and tight junction disruption alongside transcriptional changes suggestive of EndoMT (53). EndoMT is characterized by endothelial dedifferentiation towards a mesenchymal fate, involving the loss of endothelial markers and characteristics, such as key junctional architecture, and hence can result in vascular leakage (51). This pathway has been implicated in the etiology of multiple sclerosis (MS) (54) where peripheral immune activation drives BMEC transcriptional alterations which are accompanied by F-actin cytoskeletal changes akin to the phenotypes observed in our experimental models (55, 56). Crucially, peripheral inflammation is a canonical inducer of endothelial remodeling across multiple disease contexts (51) and here we implicate inflammatory signaling as a key contributor to BBB disruption and loss of endothelial identity in the allo-HSCT setting. While the neurovascular interface ordinarily protects the CNS from peripheral insults, our clinical and *in vitro* findings demonstrate that the BBB is profoundly compromised by peripheral factors characteristic of the pediatric allo-HSCT milieu, thereby limiting its capacity to protect the CNS in this context. Therefore, early and effective management of peripheral inflammation may be crucial to improve neurological outcomes.

The contribution of gastrointestinal injury to peripheral inflammation cannot be overlooked, with conditioning chemotherapy-induced gut mucosal barrier injury (i.e., gastrointestinal mucositis) considered the primary source of peripheral inflammation during HSCT (57). We identified baseline gastrointestinal vulnerabilities to poor neurological outcomes in our clinical cohort, implicating dysfunction of gut-brain axis signaling early in the timeline of pediatric allo-HSCT. Gastrointestinal injury allows translocation of gut microbiota and products (such as LPS) to systemic circulation activating innate immune cells and initiating inflammatory signaling. Accordingly, gastrointestinal mucositis is linked to the etiology of other immune mediated allo-HSCT toxicities (58). In our study, low plasma citrulline (i.e., indicative of gastrointestinal injury) was predictive of BBB damage clinically; however, *in vitro* low vs. high citrulline patient plasma did not induce significantly greater BMEC toxicity, suggesting intermediary signaling. IL-6 may act as this mediator, IL-6 concentrations temporally correlated with S100β levels, and IL-6 inhibition with tocilizumab was sufficient to attenuate allo-HSCT-induced BMEC apoptosis. Elevated IL-6 is also consistently linked to neurological deficits induced by chemotherapy (59–61) and has also been linked to other allo-HSCT toxicities (62–64), including liver endothelial cell toxicity (65).

Our findings also implicate microbiota-derived metabolites as a potentially protective component of this emerging gut-immune-BBB axis. Lower pre-transplant propionate was associated with subsequent BDNF depletion, while propionate supplementation prevented BMEC apoptosis induced by both activated immune-cell conditioned media and allo-HSCT patient plasma. Propionate and other SCFAs exert broad immunoregulatory effects, expanding regulatory T-cell populations (66, 67) and suppressing activation of CD8+ T cells (68). Accordingly, increased abundance of SCFA-producing genera prior to allo-HSCT has been associated with improved survival and reduced severity of alloreactive immune responses in pediatric recipients (69, 70), whereas reduced fecal SCFA concentrations have been linked to gut inflammation (71). Although no prior investigation has been performed investigating the influence of propionate or SCFAs on the neurological side-effects of allo-HSCT, a dietary high fiber intervention *in vivo* proved protective against chemotherapy-induced neuroinflammation, which was associated with elevated propionate production (72).

### Limitations of the study

This study is limited by the absence of neurocognitive assessment to complement the clinical biomarker analysis. Although BDNF is emerging as a marker of cognitive impairment following chemotherapy (35, 36), it incompletely captures the complexity of cognitive function, precluding direct associations between biomarkers and clinical phenotypes. The lack of patient-matched fecal samples prevented direct characterization of gut microbiota composition and limited attribution of specific taxa to phenotypes of interest. However, microbial metabolite profiling reflects microbiota functionality, i.e., higher SCFA concentrations indicative of a functionally beneficial microbiota (73–75), and certainly provide insight into microbiota-gut-brain axis interactions. The modest sample size of the clinical cohort limits generalizability, underscoring the need for validation in an independent cohort to reduce the risk of residual confounding in biomarker analyses. On the same basis, the results of our exploratory feature prioritization of baseline factors associated with post-transplant outcomes should be interpreted cautiously due to potential instability/overfitting. Similarly, the limited sample size for bulk RNA sequencing restricted its use to supporting phenotypic observations rather than enabling independent discovery. Finally, although iPSC derivation is the most functionally relevant source of BMECs (76), their use as a monoculture excludes contributions from other cells of the neurovascular unit. However, monocultured iPSC-derived BMECs present functional read-outs comparable to human *in vivo* physiology (20), and were sufficient to recapitulate key clinical phenotypes, including patient plasma-induced BMEC toxicity corresponding to elevated S100β concentrations.

## Conclusions

Our findings identify BBB dysfunction as a previously unrecognized consequence of pediatric allo-HSCT and establish the gut-immune-BBB axis as a potential driver of neurovascular injury. By integrating clinical biomarkers with mechanistic modeling, we show that gastrointestinal mucosal injury and systemic immune activation are linked to BBB dysfunction, while allo-HSCT patient plasma directly induces BMEC apoptosis *in vitro*. Managing peripheral inflammation may be crucial to prevent BBB dysfunction given radical alterations to endothelial cell phenotypes which were persistent despite concurrent anti-inflammatory intervention. Together, these findings provide a mechanistic rationale for CNS vulnerability following allo-HSCT and establish preservation of gut-immune homeostasis as a potential strategy to mitigate neurovascular side effects.

## Materials and Methods

### Materials

All reagents and materials used are listed in key materials table (Table 2).

**Table 2:**
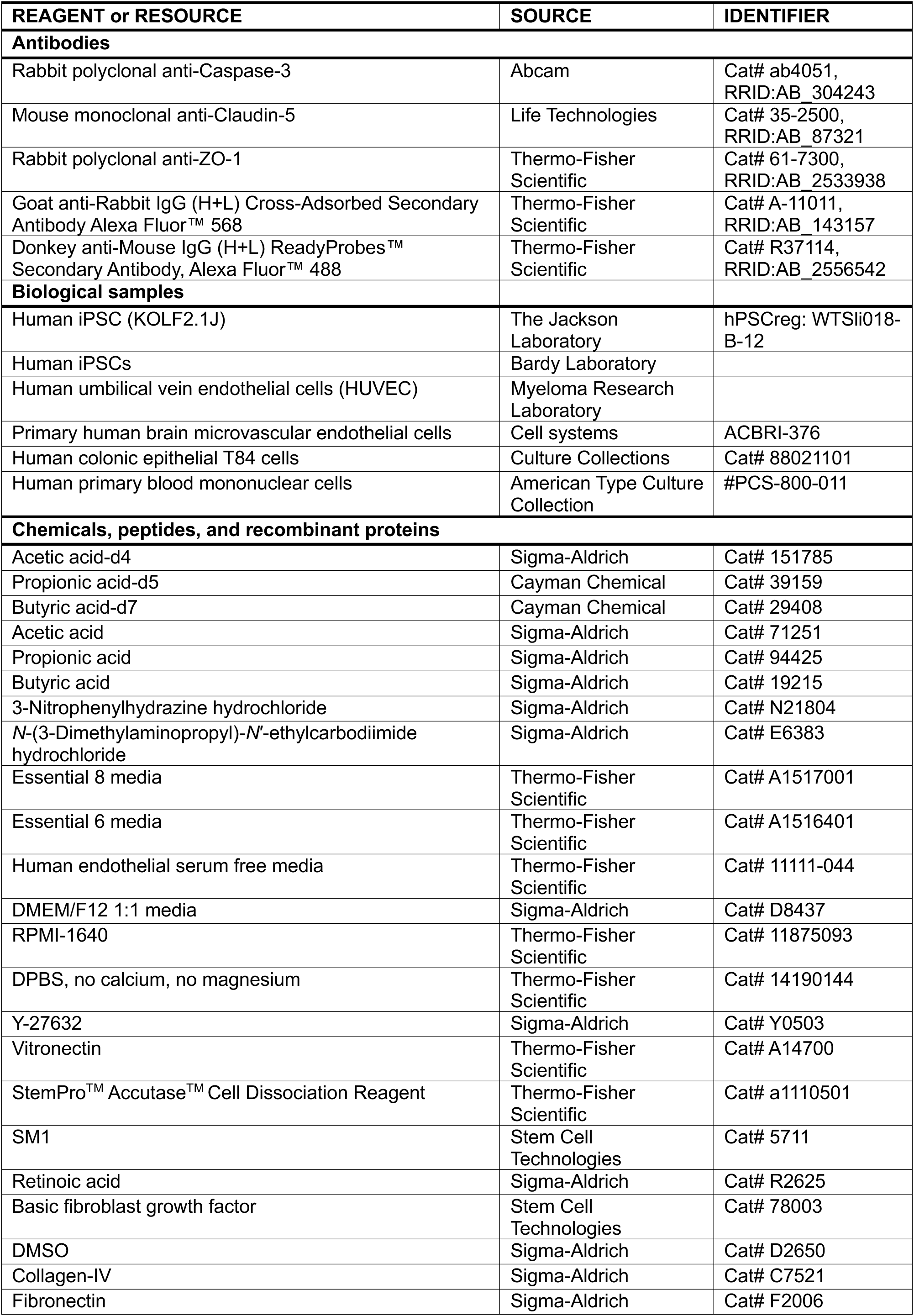
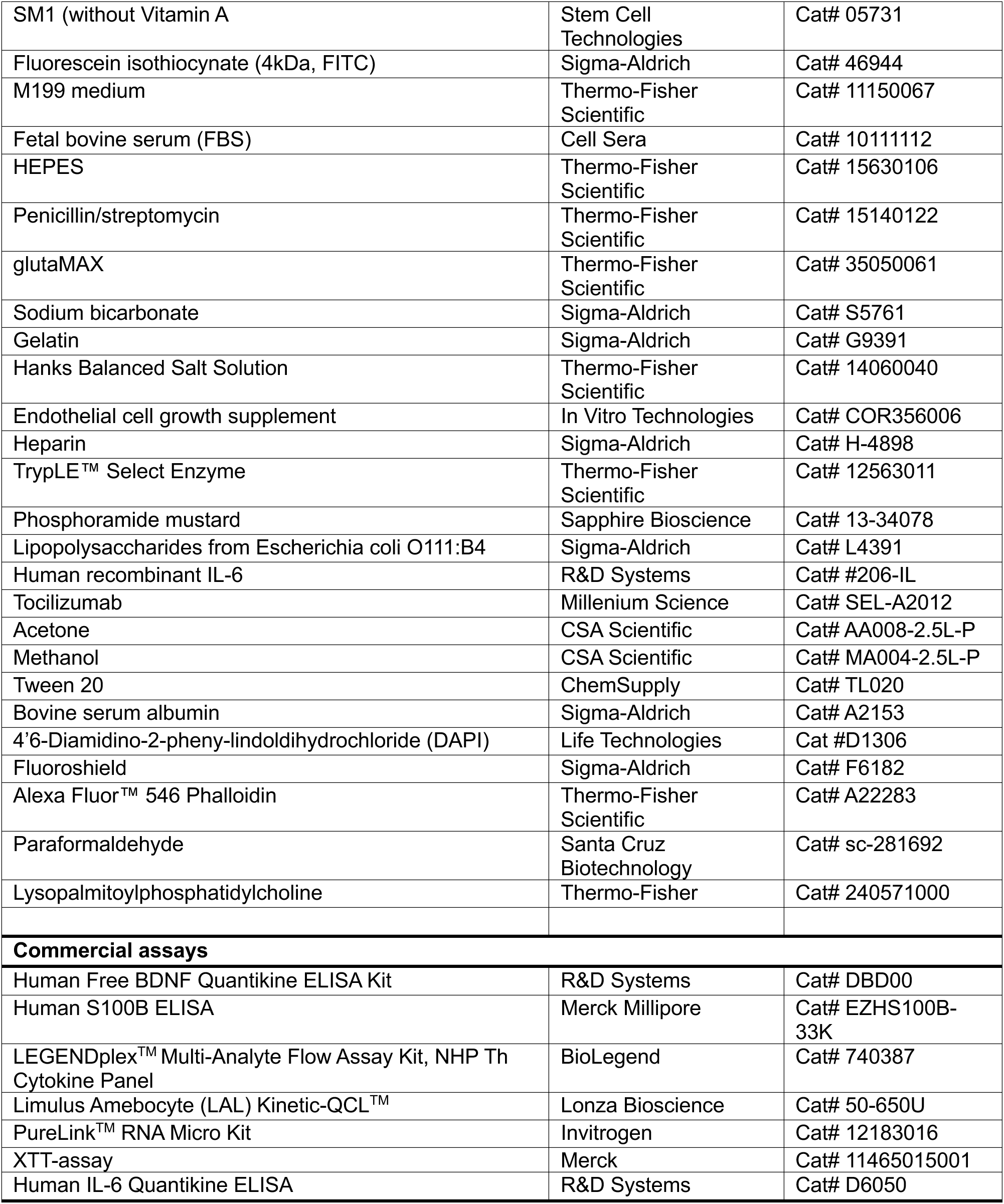
Key resources table.

### Sample collection

A research proposal was submitted to a commercial biobank (CRYOSTEM, France), the proposal was reviewed by CRYOSTEM’s Scientific Committee and approved (project no. CS-23-02), enabling access to 98 plasma samples obtained from N=47 allogeneic HSCT recipients <18-years-of age at time of transplant. A baseline, pre-HSCT (post-conditioning, but not cancer treatment naïve) sample was sourced from each patient, and a post-HSCT sample (time from HSCT highly variable, 16-391 days-post transplant), with additional follow-up samples for some patients (Fig.1A). All plasma samples were provided with a sample ID matched to a de-identified patient ID and matched demographic and clinical data.

### Enzyme-linked immunosorbent assay (ELISA)

Levels of BDNF and S100β were assayed via ELISA and samples were diluted 1:5 or used neat for BDNF and S100β, respectively. Absorbance at 450 nm was read on a BioTek Synergy HTX Multimode Reader (Agilent), and concentrations were interpolated from standard curves using GraphPad Prism v10.4.0. As ELISA guidelines specified a minimum detectable BDNF concentration <0.02ng/mL, samples with concentrations below the limit of detection (LOD) were assigned a value 5-fold below the assay sensitivity (0.004ng/mL) for analytical purposes. Sensitivity analysis confirmed that conclusions held for LOD/2 and LOD/√2.

### Multi-analyte cytokine panel

Circulating cytokines were quantified in patient samples using the LEGENDplex™ Multi-Analyte Flow Assay Kit, NHP Th Cytokine Panel, following manufacturers guidelines. Samples were diluted 1:2 and concentrations determined using a BD FACSCanto II (SAHMRI Australian Cancer Research Foundation Cellular Imaging and Cytometry Core Facility). Specifically, mean fluorescence intensity (MFI) for each analyte was measured for each standard concentration to plot a standard curve (hyperbola [X is a concentration], GraphPad Prism). MFI values for each sample were then used to interpolate concentrations of each cytokine within the plasma sample.

### Mass spectrometry

#### Plasma citrulline

Plasma citrulline was quantified at the Mass Spectrometry Imaging Core Facility (South Australian Health and Medical Research Institute, Adelaide, Australia). 10µL of each sample was combined with 25µM internal standard (citrulline-d7) and protein precipitated with 100µL methanol. Samples were centrifuged at 13,000 rpm, and the supernatant transferred to a 96-well V-bottom microtiter plate for analysis. The analysis was performed on an Agilent Infinity 1290 ultra-high performance liquid chromatography (UHPLC) using Waters ACQUITY UPLC Amide Premier (2.1×100 mm, 1.8 µm) maintained at 40°C, with a flow rate of 400µL/min. The injection volume was 2µL. Mobile phases consisted of 10mM ammonium formate in water with 0.15% formic acid (solvent A) and 10mM ammonium formate in 95% acetonitrile with 0.15% formic acid (solvent B). The samples were eluted using the following conditions: 90% B (0-1.5 min), 90%-75% B (1.5-6 min), 75%-72% B (6-8 min), 72%-50% B (8-8.1 min), 50% B (8.1-10 min), 50%-90% B (10-10.1 min), and 90% B (10.1-14 min). Mass spectrometry was performed on an API 5500 QqQ (Sciex) with the following settings: positive mode; source voltage 5500 V, curtain gas 20psi, temperature 450 °C, nebulizer 50psi, desolvation 50psi, and collisionally activated dissociation gas 9 eV. The MRM transitions were: citrulline (quant): 176.1 > 70, citrulline (qual): 176.1 > 159.1, citrulline-d7 (quant): 183.2 > 77, and citrulline-d7 (qual): 183.2 > 166.2.

#### SCFA analysis

Analysis of SCFAs in plasma samples was performed according to methods developed by Dei Cas et al (77). Deuterated internal standard master mix was diluted in 50% ACN (500µM acetate-d4, 50μM propionate-d5, 50μM butyrate-d7). External standard master mix were serially diluted in 50% ACN for a total of 6 standards. Samples and standards were combined with isopropanol then centrifuged at 21,300 x g, 15 minutes, 4°C. Supernatant from each sample or standard was collected, and combined with internal standard master, and 50mM 3-nitrophenylhydrazine (NPH), 50mM EDC (carbodiimide) and 7% pyridine in methanol. Samples, standards and blanks were derivatized at 37°C for 30 minutes and resulting solutions diluted 1:2 in 0.5% hydrochloric acid.

The derivatized samples and standards were analyzed by Agilent Infinity 1290 UHPLC coupled to Agilent 6495C Triple Quadrupole (QQQ) mass spectrometer (Agilent Technologies, Santa Clara, CA, U.S.A.). Two µL of samples with deuterated internal standards, were injected into the UHPLC-QQQ. The HPLC system was fitted with an Agilent Zorbax Eclipse C18 column (100 mm x 2.1mm x 1.8 µm) and the mobile phases consisted of 0.1% formic acid in H2O (solvent A) and 90% acetonitrile in 0.1% formic acid (solvent B). The 3-NPH derivatives of SCFAs present in the samples were eluted using the following conditions: 2%~10% B (0-7 min), 10%-60% B (7-18 min), 60%-100% B (18-19 min), 100% B (19-21 min), 100%-2% B (21-22 min), and 2% B (22-25 min). The flow rate used for elution was 300µL/min, and the column temperature was maintained at 40°C. The source parameters of the mass spectrometer were as follows: ESI interface; negative mode; Multiple reaction monitoring (MRM) scan; nebulizer 35 psi; gas temperature 220°C; gas flow rate 18L/min; sheath gas temperature, 300°C; sheath gas flow rate, 12L/min; capillary voltage, 3000 V. The MRM transition of 3-NPH derivates of SCFAs were: Acetic-NPH: 194 > 137, Acetic-d4-NPH: 194 > 137, Propionic-NPH: 208 >137, Propionic-d5-NPH: 213 > 137, Butyric-NPH: 222 > 137, Butyric-d7-NPH: 229 > 137m/z. MRM fragment voltage and collision energy were 166 V and 25 V for all derivates. The quantitative analysis was performed using Agilent MassHunter Quantitative Analysis (Quant-My-Way, version 10.1).

### Endotoxin assay

Levels of lipopolysaccharide (LPS, gram-negative bacterial endotoxin) within plasma samples were assayed using a commercially available test. Bacterial endotoxin catalyzes activation of a pro-enzyme, which results in production of p-nitroaniline (pNA) from a substrate (Ac-Ile-Glu-Ala-Arg-pNA), which produces a yellow color that can be quantified by colorimetric analysis. pNA is measured at 405nm continuously throughout the assay incubation. As endotoxin (LPS) is the catalyst for this reaction, the Sample Reaction Time, is inversely related to the endotoxin content of the sample. Hence, Total Reaction Time minus Sample Reaction Time provides semi-quantifiable insight to LPS content within the plasma sample. Absorbance throughout the reaction time was monitored using a BioTek Synergy HTX Multimode Reader (Agilent), every 150 seconds for a total of 40 reads.

### Cell culture

#### iPSC differentiation to BMECs

Human induced pluripotent stem cells (iPSCs) (either reprogrammed from fibroblasts of healthy donors in the laboratory of Prof. Bardy; passage 20-30, or KOLF2.1J cell line; passage 9) were cultured and differentiated based on methods described by Hollmann et al (78). Cells were thawed in Essential 8 (E8) media with 10μM Y-27632 and seeded onto vitronectin-coated wells of a 6-well plate. 24-hours post-thaw, E8 media was refreshed without Y-27632. Once 70% confluence was reached, single-cell suspension was achieved via incubation with StemPro™ Accutase™ Cell Dissociation Reagent (Accutase) for 5 minutes at 37°C, and counted via Trypan Blue staining and a Countess™ 3 FL Automated Cell Counter (Invitrogen, #AMQAF2000). Cells were then seeded at 125,000 cells per well of vitronectin-coated 6-well plate in E8 with 10μM Y-27632. 24-hours later media was replaced with Essential 6 (E6) media to initiate differentiation (D0).

E6 media was refreshed daily for 4 days and then media was replaced with human endothelial serum free media (HESFM) with 1X SM1, 10μM retinoic acid (RA,) and 20ng/mL basic fibroblast growth factor (bFGF) for BMEC expansion – media was not changed for 48-hours. On D6 cells were dissociated via 20-minute Accutase incubation (37°C) and then collected via centrifuge. Cells were resuspended in HESFM + SM1 and counted, then 10% DMSO was added to cryopreserve BMEC stocks.

BMEC stocks were rapidly thawed as needed and resuspended in HESFM media +SM1, RA, bFGF and 10μM Y-27632 then and seeded at 100,000 cells/cm^2^ on collagen-IV/fibronectin coated inserts of 24-well transwell plates (0.4μm pore size, polyester membrane, Corning, #CLS3470). Extracellular matrix coating solution was composed of a 5:4:1 coating solution of deionized water, 1mg/mL collagen-IV, 1mg/mL fibronectin, incubated at 37°C for a minimum of 4-hours, up to overnight. 24-hours post thaw, media was refreshed with HESFM +SM1 (without Vitamin A), and at 48-hours cells were either stained and fixed for phenotyping, collected for RNA extraction, or exposed to treatment conditions. To assess barrier functionality permeability of 4kDa fluorescein isothiocyanate (FITC) from the apical to basolateral compartment was also assayed. FITC was reconstituted in DPBS at 10mg/mL FITC and added to apical transwell compartment culture media at a final concentration of 200μg/mL. For model characterization, the basolateral compartment was sampled at 0, 3, 6, 24 and 48-hours, and for experimental conditions sampling was performed at 24 and 48-hours. Fluorescence of basolateral media was quantified on a BioTek Synergy HTX Multimode Reader (Agilent) and concentrations interpolated from fluorescence measures of known standards. Trans-endothelial electrical resistance (TEER) measurements were also taken after 48-hours on transwells using the EVOM Manual™ with STX-HTS electrodes (World Precision Instruments) to determine barrier functionality.

#### HUVEC cultures

Cryopreserved human umbilical vein endothelial cells (HUVECs) (kind gift from Myeloma Research Laboratory) were thawed and maintained in M199 medium with additives: 10-20% fetal bovine serum (FBS), HEPES, 100IU/mL penicillin and 100μg/mL streptomycin (P/S), and 1X glutaMAX, on 2% gelatin-coated (2% w/v gelatin dissolved in deionized water with FBS, sodium bicarbonate, Hanks Balanced Salt Solution) T75 culture flasks. Growth media was supplemented with 30μg/mL endothelial cell growth supplement (ECGS) and 15μg/mL heparin immediately prior to refresh. HUVECs were passaged with Accutase, undergoing a maximum of 15 passages. HUVECs were sub-cultured at 100,000 cells/cm^2^ on 2% gelatin-coated transwells, allowed to adhere 24-hours before media refresh, before being exposed to treatment conditions 48-hours post-subculture.

#### Primary BMEC cultures

Primary human brain microvascular endothelial cells (pBMECs, passage 3) were sourced commercially and cultured in DMEM/F12 1:1 media with 10% FBS, 1X glutaMAX, P/S and 40μg/mL ECGS in 6-well plates coated with either 2% gelatin-mixture or collagen-IV (400μg/mL) and fibronectin (100μg/mL) extracellular matrix in deionized water. pBMEC seeding density was 125,000 cells/well.

#### Colonic epithelial cultures

Human colonic epithelial T84 cells were cultured in DMEM/F12 Ham 1:1 media with additives of 10% FBS, glutaMAX, and P/S in T75 flasks. T84s were passaged using TrypLE™ Select Enzyme and maintained between 5-15 passages. For PM dose-finding study, T84s were seeded at 100,000 cells/cm^2^ in 96-well plate, 24-hours prior to treatment. For *in vitro* gut permeability assessment, T84s were seeded at 100,000 cells/cm^2^ onto transwell inserts of a 24-well transwell plate and allowed 4-5 days to reach physiological TEER values, prior to addition of PM to culture media.

#### Primary PBMC culture

Normal human primary blood mononuclear cells (PBMCs) were sourced commercially and cultured at 5×10^6^ cells/mL in RPMI-1640 media containing 10% FBS, P/S and glutaMAX in T25 flasks.

### RNA extraction, sequencing and analysis

For RNA extractions to investigate cellular identity, all cells (excluding pBMECs) were seeded at 750,00 cells/cm^2^ on 24-well transwell plate inserts, with a minimum of 5 transwells pooled per replicate for a total N=3, and cultured for 48-hours prior to collection. For pBMECs, cells were thawed at 125,000 cells/cm^2^ in 6-well plates (2 wells pooled per replicate, final N=2) and cultured for 24-hours prior to collection. For RNA extractions following treatment conditions, BMECs were initially seeded at 100,000 cells/cm^2^ on 24-well transwell plate inserts and cultured for 48-hours prior to treatment initiation, with treatment exposures 48-hours in duration. Each treatment condition had a minimum of 5 transwells per replicate which were pooled into 2 replicates for sequencing.

RNA extractions were performed using the PureLink™ RNA Micro Kit following manufacturer’s instructions. Briefly, cells were lysed with Lysis Buffer (+1mM Dithiothreitol [DTT] and 70pg/mL Carrier RNA) and transferred to collection tubes. For cells cultured on transwell inserts, inserts were also cut out using a scalpel blade and added to collection tubes. Lysates were then homogenized via repeated passing through a 21-gauge needle, washed with 70% ethanol, and vortexed to dissolve precipitate. Samples were processed through a PureLink™ Micro Kit Column and washed with washed buffer before incubation with PureLink™ DNase for 15 minutes at room temperature. Wash steps performed as specified in manufacturer’s guidelines and then RNA was eluted in RNase-Free Water. Subsequent RNA concentrations and purity were determined via a NanoDrop 8000 Spectrophotometer (Thermo-Fisher).

RNA sequencing and major analysis was performed by the South Australian Genomics Centre (South Australian Health and Medical Research Institute, Adelaide, Australia). mRNA libraries were prepared using the Nugen Universal Plus mRNA-seq protocol, including 12 cycles of amplification. Equimolar pools were prepared and converted to MGI compatible libraries using the MGI conversion kit. Library sequencing was performed using a DNBSEQ-G400 Flow Cell PE100 (MGI, Shenzhen, China). RNA-seq data processing was performed using the nf-core/rnaseq pipeline (v3.19.0, doi: 10.5281/zenodo.1400710), which generated Salmon (v1.10.3) count tables for transcript quantification. Differential expression analysis was conducted using the nf-core/differentialabundance pipeline (v1.5.0, doi: 10.5281/zenodo.7568000), with DESeq2 (v1.34.0) used to identify differentially expressed genes. Gene set enrichment analysis was performed using gprofiler2 with the GO Biological Process (GO:BP) database as the annotation source and multiple testing correction was applied with the g:SCS method. Redundant GO terms were subsequently reduced by semantic similarity and grouped under representative parent terms using the rrvgo package. The top 20 differentially enriched parent terms, ranked by differential index, were visualized as bar charts. Dotplots were generated to visualize comparison of BMEC gene expression to implicated transcription factors and mesenchymal markers published in the literature (50, 51). 0 q-values were replaced with a minimal positive value (1×10^−300^) to allow all genes to be represented.

### Experiments

#### T84 PM dose optimization

T84 cells were seeded at 100,000 cells/cm^2^ in a 96-well plate for preliminary dose-viability study. PM was reconstituted in PBS and T84s were treated with doses (0, 0.02, 0.04, 0.08, 0.16, 0.32, 0.64, 1.28mM) for 24 and 48-hours (N=6 wells per concentration) prior to viability assessment via XTT-assay, performed via manufacturers guidelines. Briefly, after drug incubation XTT labelling mixture (0.3mg/mL) was added to each well and incubated for 4-hours at 37°C, before spectrophotometrical absorbance (450-500nm) was determined using a GloMax® Discover Microplate Reader (Promega, Australia). From this dose-finding viability study, T84s were then cultured on transwell inserts (seeded at 100,000 cells/cm^2^, N=4 wells/condition) and TEER measures were recorded daily until reaching physiological values (>1000Ω), media in the apical compartment was then refreshed with 0 (volume-matched vehicle [PBS] control), 0.32mM, 0.64 or 1.28mM PM. TEER recordings were taken at 24 and 48-hours, and at 48-hours cells were also fixed for immunocytochemistry (ICC).

#### BMEC exposure to candidates of peripheral-to-central inflammation delivery

For all experiments using iPSC-derived BMECs, a minimum of 3 transwells were allocated per condition (unless otherwise specified) per experiment and each experiment performed twice (N≥6 wells). BMECs were cultured on transwell inserts and molecular/drug candidates of peripheral-to-central delivery of toxicity were added to apical compartment culture media to characterize their capacity to induce barrier damage. Specifically, cells were exposed to either 10ng/mL Lipopolysaccharides from Escherichia coli O111:B4 (LPS), 1.28mM PM or a media only control, for 48-hours. BMECs were also exposed to either 10ng/mL human recombinant IL-6 or 10ng/mL IL-6 + 50μg/mL of the humanized monoclonal antibody tocilizumab (TCZ, anti-IL-6 receptor) and this was compared to both a media only control and media + 50μg/mL TCZ. Sampling of basolateral compartment for FITC permeability was performed at 24- and 48-hours and after 48-hours, cells were fixed and stained for outcomes of interest. RNA was also extracted from PM-treated BMECs for RNA-seq.

#### Mimicking cancer-treatment induced peripheral inflammation in vitro

10ng/mL LPS and 1.28mM PM (N=3) or matched volume PBS (N=1) were added to the culture media of primary PBMCs cultured in T25 flasks for 24-hours. After this stimulation period, PBMC supernatant was collected, and the number of live cells were counted via Trypan blue staining and Countess Automated Cell Counter to determine viability. IL-6 levels within the supernatant were used as a surrogate marker of peripheral inflammation. PBMC supernatant was diluted 1:50 and IL-6 concentrations were quantified via Human IL-6 Quantikine ELISA performed as per manufacture’s guidelines on samples run in duplicate.

#### In vitro BBB stimulation with PBMC conditioned media and testing of potential interventions

PBMC conditioned media (LPS+PM: stim-PBMC-CM; or PBS: ctrl-PBMC-CM) was diluted 50% in BMEC culture media and added to the apical transwell compartment for 48-hours. To ensure that observed effects were not a result of exposure to PBMC culture media, depletion of normal endothelial media, or the secretions of unstimulated PBMCs, a normal cell media only control was also included. Given no significant differences between media only control or ctrl-PBMC-CM for all outcomes of interest, and given PBMC cultures were preferentially stimulated with stim-PBMC-CM to ensure sufficient supernatant for all experiments, a media only control was used for the remainder of all experiments. In addition to stim-PBMC-CM, interventions of interest were applied in parallel. Specifically, either TCZ or propionate were added to either BMEC culture media, or BMEC media + 50% stim-PBMC-CM. Final TCZ concentration was 50μg/mL and final concentration of propionate was 1μM. BMECs were exposed to PBMC CM ± interventions for 48-hours.

#### Testing barrier damaging effects of human plasma from pediatric allogeneic HSCT recipients

Plasma from post-HSCT sampling time points were sorted by plasma citrulline concentration (clinical biomarker of gut mucosal barrier integrity) into 5 samples with highest citrulline concentrations and 5 samples with lowest concentrations. These samples were then pooled and diluted 25% into BMEC media (+P/S) and BMECs were exposed to 25% plasma from pediatric patients (either high or low citrulline concentrations) post-HSCT for 24-hours. Given that BMECs are usually cultured without antibiotics, 1X P/S was also added to BMEC media to reduce risk of contamination, with media +antibiotics (media+Ab) used as the matched control. To ensure that observed effects were not only a result of exposing cells to human plasma, in parallel, BMECs were also exposed 25% plasma from a healthy adult control (HREC project code: H-2024-117). Interventions (50μg/mL TCZ or 1μM propionate) were diluted in media+ patient plasma mixtures in parallel to plasma only (25%) treatment. Due to limited availability of patient plasma samples, each experiment had ≥1 transwell per condition, with a total of 2 individual experiments (N=2 biological replicates).

### Immunocytochemistry

To fix cells in transwells, media was removed and apical and basolateral compartments washed with ice-cold DPBS to remove cell culture media. The fixing solution of 1:1 (v/v) acetone and methanol at −20°C was then added to the apical compartment and incubated for 15 minutes at room temperature. Fixing solution was then aspirated and cells washed with PBS prior to 3-minute permeabilization with 0.1% Tween 20 in PBS. After the TX-100 was completely removed with PBS wash, cells were blocked for 1-hour at room temperature in 4% bovine serum albumin (BSA), then incubated with primary antibodies (anti-caspase-3 1:1000; anti-ZO-1 1:1000; anti-claudin-5 1:50) diluted in 1% BSA – either for 1-hour at room temperature or overnight at 4°C. After incubation, primary antibody was removed thoroughly with 3x PBS washes, followed by 1-hour incubation at room temperature with secondary antibodies (anti-rabbit Alexa Fluor™ 568; anti-mouse Alexa Fluor™) diluted 1:250 in 1% BSA. Secondary antibodies were removed via aspiration and PBS wash and then cell nuclei were stained with 1μg/mL 4’6-Diamidino-2-pheny-lindoldihydrochloride (DAPI) for 10 minutes, also at room temperature. After DAPI aspiration and PBS wash, transwell inserts were cut out using a scalpel and mounted onto microscope slides in Fluoroshield mounting medium.

#### F-actin staining with phalloidin

To stain F-actin cytoskeleton fibers with phalloidin, a simultaneous fixing, permeabilization and staining solution was made-up via diluting 6.6μM Alexa Fluor™ 546 Phalloidin 1:400 in 4% paraformaldehyde (PFA) containing 100μg/mL lysopalmitoylphosphatidylcholine. Cell culture media was removed and apical and basolateral transwell compartments washed with ice-cold DPBS. The phalloidin staining solution was added to the apical compartment and incubated for 20 minutes at 4°C. The staining solution was aspirated and washed with PBS, before addition of 1μg/mL DAPI for a 10-minute incubation at room temperature. After nuclei staining, transwell inserts were cut out using a scalpel blade and mounted onto coverslips in Fluoroshield for imaging.

### Imaging and analysis

Cells were visualized using a Confocal Olympus FV300 microscope (Adelaide Microscopy, Australia) using a 100X immersion-oil objective. DAPI staining was used to initially focus the microscope and then 3 DAPI+ fields of view (FOVs) were selected at random across each stained well for each condition (minimum of 6 FOVs for all stains). Capase-3 staining was imaged at 1X magnification and for high resolution membrane staining (ZO-1, CD144, Claudin-5) and F-actin staining, a 2X and 1.5X magnification was applied, respectively. To quantify immunofluorescent and phalloidin staining, .tiff images per experiment were imported into ImageJ2 version 2.16.

For caspase-3 staining all.tiff files stacks were imported as an image sequence split into color channels, with the red channel used for further analysis. A threshold was applied universally to all images using the ‘moments’ method and %area stain values were measured. FOVs for which background impacted the accuracy of the threshold method were excluded manually.

Cell size was quantified using ZO-1 staining. All images from the same independent experiment were imported as an image sequence. In each field of view (FOV), three cells were manually outlined using the membrane-associated ZO-1 signal to define cell boundaries, and the area was calculated. When fewer than three cells per FOV had fully visible borders, the maximal number of cells with complete boundaries were measured. Mean cell area was calculated per well from the three FOVs, and the average cell area of the corresponding control condition was determined. Cell areas for all conditions were then expressed as a percentage of the mean control cell area.

To analyze tight junction structure using ZO-1 staining, methods were based on the approach developed by Ebert et al (79). .tiff files were imported as an image sequence, a uniform scale was applied, and also a universal threshold was applied across all images using the auto-threshold method. A 10μm straight line was drawn longitudinally along the junctional ZO-1 staining and the staining intensity along the line was plotted within ImageJ using the ‘Analyze’ > ‘Plot profile’ function: plotting distance (in μm) along the line on the x-axis and fluorescence intensity on the y-axis. Numeric values were exported using the ‘List’ function. This analysis was performed for three junctions per FOV. Extracted data were imported into GraphPad Prism (v10.5.0). Two datasets were generated: one containing measurements from each individual line and another in which the nine measurements per condition (3 junctions across 3 FOVs) were averaged for visualization. Using the individual-line dataset, linear regression was performed for each junction and the standard deviation of the residuals (sy.x) was calculated to quantify the average deviation of data points from the regression line. The sy.x value was used as an indicator of irregularity in junctional ZO-1 staining. Sy.x values were averaged per FOV and grouped by treatment condition for statistical analysis.

### Statistical analysis

All data analyses were performed using GraphPad Prism (version 10.4.0) unless otherwise specified, and specific statistical tests are detailed in the associated figure legends. For patient data, descriptive statistics (median, IQR) were used to summarize trends in the patient characteristics and to define threshold scores for decreased BDNF (BDNF<1^st^ IQR) and BBB damage (S100β>4^th^ IQR). Data was initially tested for normality via the Shapiro-Wilk test, with a statistical decision rule applied: if all variables for a biomarker passed normality, a parametric test was applied otherwise a non-parametric test was performed. Outliers were identified using the ROUT method (Q=1%). A student’s t-test (parametric) or Mann-Whitney test (non-parametric) were used to identify significant differences between pre- and post-HSCT. Unpaired tests were selected because some participants had multiple post-HSCT samples, and the primary goal was to assess overall group-level differences rather than inter-individual changes. For multiple post-HSCT samples for the same participant, averages were also calculated to confirm that results were unchanged when accounting for participant-level clustering (data not shown). Student’s unpaired t-tests (for parametric data) or Mann-Whitney tests (for non-parametric data) were also performed to identify significant differences between GvHD- and GvHD+ samples post-HSCT.

For longitudinal characterization samples were binned into 50-day intervals: 0=pre-HSCT, 1-50-days-post-HSCT, 51-100-days-post-HSCT, 101-150-days-post-HSCT. Given limited samples (N=7) for >150-days-post-HSCT, longitudinal analysis was restricted to <150-days. Mixed-effects analysis was used to determine statistically significant within-marker changes between baseline and the binned time-points post-HSCT for parametric data, with a Kruskal-Wallis test performed for non-parametric data. Longitudinal BDNF analyses were repeated in Stata to confirm results and to verify that age-at-diagnosis did not confound outcomes. Correlation matrices were generated in Python using the ‘pearsonr’ function of the scipy.stats module to explore relationships among biomarkers and to identify highly correlated measures. These analyses were exploratory and intended to inform interpretation rather than to support inclusion of multiple biomarkers in the same regression model. To avoid the potential for multicollinearity, biomarkers were not included simultaneously in multivariable longitudinal models. For survival analysis, median post-HSCT biomarker values were calculated, and samples were stratified into “low” or “high” groups relative to the median. Kaplan-Meier survival curves were plotted, with significance tested using the Gehan-Breslow-Wilcoxon test. Univariable Cox regression was used to estimate hazard ratios and 95% confidence intervals; multivariable models were not performed due to limited sample size and events.

Exploratory analyses of baseline factors and neurological outcomes were performed in Python (version 3.10.4). Non-numerical values were first encoded to the binary, and variables with >50% data missing (e.g., acute GvHD grade) were excluded. For patients with multiple post-HSCT samples, the most extreme value (i.e., lowest BDNF value, highest S100β value) was adopted to understand the maximum risk of the patient in the ‘worst-case’ scenario. Pearson correlation coefficients and P-values were computed using the ‘pearsonr’ function from the scipy.stats module to identify baseline factors (prior to HSCT) most associated with (1) low plasma BDNF (1^st^ IQR), or (2) BBB damage (4^th^ IQR), post-HSCT.

For *in vitro* experiments, normality was assessed using the Shapiro-Wilk test and potential outliers were tested via the ROUT method (Q=1%). If all variables passed normality, unpaired t-test or one-way-ANOVA were performed, otherwise Mann-Whitney, or Kruskal-Wallis tests were performed for non-parametric data. For repeated measures of FITC-permeability, mixed-effects analysis was used. For dose-response assessment, simple linear regression analysis was performed with PM concentration as the predictor and cell viability as the outcome. Statistical significance was defined as P<0.05 for all analyses.

### Data and code availability

RNA sequencing data generated in this study will be made publicly available on the NCBI database under the BioProject accession number PRJNA1444808 upon publication.

## Acknowledgements

The authors would like to acknowledge CRYOSTEM for access to the biobank, and the patients and their families for access to the clinical samples. We would like to thank Ms. Bronwyn Cambareri for her advice and technical support, Mr. Cheng Lu for his contributions to the statistical analysis, and Dr. John Salamon from the South Australian Genomics Centre for his work on the bioinformatics analysis. Plasma citrulline analysis was performed at the Mass Spectrometry Imaging Core Facility (South Australian Health and Medical Research Institute, Adelaide, Australia) and RNA sequencing and major analysis was performed by the South Australian Genomics Centre (South Australian Health and Medical Research Institute, Adelaide, Australia).

## Author contributions

Conceptualization, MRD, CBC, HRW; investigation, MRD, CBC, YL, AL, HRW; methodology, MRD, CBC, FJR, YL, ZG, CMDW, CSB, CB, HRW; data analysis: MRD, MD, AS; funding acquisition, HRW; resources, MRD, AL, ACWZ, CB, HRW; supervision, CB, ACWZ, HRW; visualization, MRD; writing (original draft), MRD; writing (review and editing), MRD, CBC, FJR, YL, MD, ZG, AS, CMDW, AL, ACWZ, CSB, CB, HRW.

## References

1. Remberger M, Ackefors M, Berglund S, Blennow O, Dahllöf G, Dlugosz A, et al. Improved survival after allogeneic hematopoietic stem cell transplantation in recent years. A single-center study. Biol Blood Marrow Transplant. 2011;17(11):1688–97.

2. Turcotte LM, Whitton JA, Leisenring WM, Howell RM, Neglia JP, Phelan R, et al. Chronic conditions, late mortality, and health status after childhood AML: a Childhood Cancer Survivor Study report. Blood. 2023;141(1):90–101.

3. Clarke SA, Skinner R, Guest J, Darbyshire P, Cooper J, Vora A, et al. Clinical outcomes and health-related quality of life (HRQOL) following haemopoietic stem cell transplantation (HSCT) for paediatric leukaemia. Child: Care, Health and Development. 2011;37(4):571–80.

4. Olsson M, Aili K, Jarfelt M, Nygren JM, Arvidsson S. Life is an ongoing existential battle – experiences from adult survivors after allogeneic hematopoietic stem cell transplantation during childhood acute lymphoblastic leukemia. European Journal of Oncology Nursing. 2025;77:102929.

5. Visentin S, Auquier P, Bertrand Y, Baruchel A, Tabone M-D, Pochon C, et al. The Impact of Donor Type on Long-Term Health Status and Quality of Life after Allogeneic Hematopoietic Stem Cell Transplantation for Childhood Acute Leukemia: A Leucémie de l’Enfant et de L’Adolescent Study. Biology of Blood and Marrow Transplantation. 2016;22(11):2003–10.

6. Giralt S, Bishop MR. Principles and overview of allogeneic hematopoietic stem cell transplantation. Cancer Treat Res. 2009;144:1–21.

7. Lindsay J, Kerridge I, Wilcox L, Tran S, O’Brien TA, Greenwood M, et al. Infection-Related Mortality in Adults and Children Undergoing Allogeneic Hematopoietic Cell Transplantation: An Australian Registry Report. Transplant Cell Ther. 2021;27(9):798.e1–.e10.

8. Gassas A, Sivaprakasam P, Cummins M, Breslin P, Patrick K, Slatter M, et al. High transplant-related mortality associated with haematopoietic stem cell transplantation for paediatric therapy-related acute myeloid leukaemia (t-AML). A study on behalf of the United Kingdom Paediatric Blood and Bone Marrow Transplant Group. Bone Marrow Transplant. 2018;53(9):1165–9.

9. Loeffen EAH, Knops RRG, Boerhof J, Feijen E, Merks JHM, Reedijk AMJ, et al. Treatment-related mortality in children with cancer: Prevalence and risk factors. Eur J Cancer. 2019;121:113–22.

10. Berbis J, Michel G, Chastagner P, Sirvent N, Demeocq F, Plantaz D, et al. A French cohort of childhood leukemia survivors: impact of hematopoietic stem cell transplantation on health status and quality of life. Biol Blood Marrow Transplant. 2013;19(7):1065–72.

11. Armenian SH, Sun CL, Kawashima T, Arora M, Leisenring W, Sklar CA, et al. Long-term health-related outcomes in survivors of childhood cancer treated with HSCT versus conventional therapy: a report from the Bone Marrow Transplant Survivor Study (BMTSS) and Childhood Cancer Survivor Study (CCSS). Blood. 2011;118(5):1413–20.

12. Bhatia S. Long-term health impacts of hematopoietic stem cell transplantation inform recommendations for follow-up. Expert Rev Hematol. 2011;4(4):437–52; quiz 53-4.

13. Ansari S, Garg A, Khan MA. Neurocognitive outcomes in pediatric hematological cancer survivors post-HSCT: A systematic review. Clin Transplant. 2024;38(1):e15193.

14. Yen HJ, Eissa HM, Bhatt NS, Huang S, Ehrhardt MJ, Bhakta N, et al. Patient-reported outcomes in survivors of childhood hematologic malignancies with hematopoietic stem cell transplant. Blood. 2020;135(21):1847–58.

15. Wu NL, Krull KR, Cushing-Haugen KL, Ullrich NJ, Kadan-Lottick NS, Lee SJ, Chow EJ. Long-term neurocognitive and quality of life outcomes in survivors of pediatric hematopoietic cell transplant. J Cancer Surviv. 2022;16(3):696–704.

16. Gabriel M, Hoeben BAW, Uhlving HH, Zajac-Spychala O, Lawitschka A, Bresters D, Ifversen M. A Review of Acute and Long-Term Neurological Complications Following Haematopoietic Stem Cell Transplant for Paediatric Acute Lymphoblastic Leukaemia. Front Pediatr. 2021;9:774853.

17. Bleggi-Torres LF, de Medeiros BC, Werner B, Neto J, Loddo G, Pasquini R, de Medeiros CR. Neuropathological findings after bone marrow transplantation: an autopsy study of 180 cases. Bone Marrow Transplantation. 2000;25(3):301–7.

18. Kadry H, Noorani B, Cucullo L. A blood–brain barrier overview on structure, function, impairment, and biomarkers of integrity. Fluids and Barriers of the CNS. 2020;17(1):69.

19. Knox EG, Aburto MR, Clarke G, Cryan JF, O’Driscoll CM. The blood-brain barrier in aging and neurodegeneration. Molecular Psychiatry. 2022;27(6):2659–73.

20. Davies MR, Cross CB, Aburto MR, Cryan JF, Wardill HR. In vitro models of microbiota-gut-brain axis communication at the blood-brain barrier interface. Journal of Cerebral Blood Flow & Metabolism. 2026:0271678X261419964.

21. Chen T, Dai Y, Hu C, Lin Z, Wang S, Yang J, et al. Cellular and molecular mechanisms of the blood–brain barrier dysfunction in neurodegenerative diseases. Fluids and Barriers of the CNS. 2024;21(1):60.

22. Vance ML, Nagy D, Brunner E, Morkotinis V, Black JL, Refai LH, et al. Endothelial-to-mesenchymal transition in the central nervous system: A potential therapeutic target to combat age-related vascular fragility. The Journal of Pharmacology and Experimental Therapeutics. 2025;392(11):103747.

23. Park JC, Chang L, Kwon H-K, Im S-H. Beyond the gut: decoding the gut–immune–brain axis in health and disease. Cellular & Molecular Immunology. 2025;22(11):1287–312.

24. O’Riordan KJ, Moloney GM, Keane L, Clarke G, Cryan JF. The gut microbiota-immune-brain axis: Therapeutic implications. Cell Reports Medicine. 2025;6(3):101982.

25. Logsdon AF, Erickson MA, Rhea EM, Salameh TS, Banks WA. Gut reactions: How the blood-brain barrier connects the microbiome and the brain. Exp Biol Med (Maywood). 2018;243(2):159–65.

26. Li H, Sun J, Du J, Wang F, Fang R, Yu C, et al. Clostridium butyricum exerts a neuroprotective effect in a mouse model of traumatic brain injury via the gut-brain axis. Neurogastroenterology & Motility. 2018;30(5):e13260.

27. Giangiulio O, Maccarone R. The Blood–Brain Barrier as an Integration Hub in Alzheimer’s Disease: How Microbiota Metabolites Modulate Central Signal Processing. CNS Neuroscience & Therapeutics. 2025;31(12):e70703.

28. Tang C-F, Wang C-Y, Wang J-H, Wang Q-N, Li S-J, Wang H-O, et al. Short-Chain Fatty Acids Ameliorate Depressive-like Behaviors of High Fructose-Fed Mice by Rescuing Hippocampal Neurogenesis Decline and Blood–Brain Barrier Damage. Nutrients [Internet]. 2022; 14(9):[1882 p.].

29. Liu J, Jin Y, Ye Y, Tang Y, Dai S, Li M, et al. The neuroprotective effect of short chain fatty acids against sepsis-associated encephalopathy in mice. Frontiers in Immunology. 2021;12:626894.

30. Dalile B, Van Oudenhove L, Vervliet B, Verbeke K. The role of short-chain fatty acids in microbiota–gut–brain communication. Nature Reviews Gastroenterology & Hepatology. 2019;16(8):461–78.

31. Karlik JB, Kesavan A, Nieder ML, Hawks R, Jin Z, Bhatia M, Ladas EJ. Plasma citrulline as a biomarker for enterocyte integrity in pediatric blood and BMT. Bone Marrow Transplantation. 2014;49(3):449–50.

32. Herbers AH, de Haan AF, van der Velden WJ, Donnelly JP, Blijlevens NM. Mucositis not neutropenia determines bacteremia among hematopoietic stem cell transplant recipients. Transpl Infect Dis. 2014;16(2):279–85.

33. Undén J, Romner B. Can low serum levels of S100B predict normal CT findings after minor head injury in adults?: an evidence-based review and meta-analysis. J Head Trauma Rehabil. 2010;25(4):228–40.

34. Ingebrigtsen T, Romner B, Kock-Jensen C. Scandinavian guidelines for initial management of minimal, mild, and moderate head injuries. The Scandinavian Neurotrauma Committee. J Trauma. 2000;48(4):760–6.

35. Ng DQ, Chan D, Agrawal P, Zhao W, Xu X, Acharya M, Chan A. Evidence of brain-derived neurotrophic factor in ameliorating cancer-related cognitive impairment: A systematic review of human studies. Crit Rev Oncol Hematol. 2022;176:103748.

36. Ng DQ, Cheng I, Wang C, Tan CJ, Toh YL, Koh YQ, et al. Brain-derived neurotrophic factor as a biomarker in cancer-related cognitive impairment among adolescent and young adult cancer patients. Scientific Reports. 2023;13(1):16298.

37. Trudeau J, Ng DQ, Sayer M, Tan CJ, Ke Y, Chan RJ, Chan A. Brain-derived neurotrophic factor and cytokines as predictors of cognitive impairment in adolescent and young adult cancer patients receiving chemotherapy: a longitudinal study. BMC Cancer. 2025;25(1):1045.

38. Yap NY, Tan NYT, Tan CJ, Loh KW-J, Ng RCH, Ho HK, Chan A. Associations of plasma brain-derived neurotrophic factor (BDNF) and Val66Met polymorphism (rs6265) with long-term cancer-related cognitive impairment in survivors of breast cancer. Breast Cancer Research and Treatment. 2020;183(3):683–96.

39. Vikner T, Garpebring A, Björnfot C, Nyberg L, Malm J, Eklund A, Wåhlin A. Blood–brain barrier integrity is linked to cognitive function, but not to cerebral arterial pulsatility, among elderly. Scientific Reports. 2024;14(1):15338.

40. Barisano G, Montagne A, Kisler K, Schneider JA, Wardlaw JM, Zlokovic BV. Blood-brain barrier link to human cognitive impairment and Alzheimer’s Disease. Nat Cardiovasc Res. 2022;1(2):108–15.

41. Che J, Sun Y, Deng Y, Zhang J. Blood-brain barrier disruption: a culprit of cognitive decline? Fluids and Barriers of the CNS. 2024;21(1):63.

42. Alkayed NJ. Blood-Brain Barrier: A Shield Against Cognitive Decline. Stroke. 2024;55(12):2906–8.

43. Hart E, Odé Z, Derieppe MPP, Groenink L, Heymans MW, Otten R, et al. Blood-brain barrier permeability following conventional photon radiotherapy - A systematic review and meta-analysis of clinical and preclinical studies. Clin Transl Radiat Oncol. 2022;35:44–55.

44. Nakkazi A, Forster D, Whitfield GA, Dyer DP, Dickie BR. A systematic review of normal tissue neurovascular unit damage following brain irradiation-Factors affecting damage severity and timing of effects. Neurooncol Adv. 2024;6(1):vdae098.

45. Yuan H, Gaber MW, McColgan T, Naimark MD, Kiani MF, Merchant TE. Radiation-induced permeability and leukocyte adhesion in the rat blood-brain barrier: modulation with anti-ICAM-1 antibodies. Brain Res. 2003;969(1-2):59–69.

46. Lu TM, Houghton S, Magdeldin T, Durán JGB, Minotti AP, Snead A, et al. Pluripotent stem cell-derived epithelium misidentified as brain microvascular endothelium requires ETS factors to acquire vascular fate. Proc Natl Acad Sci U S A. 2021;118(8).

47. Karyekar CS, Fasano A, Raje S, Lu R, Dowling TC, Eddington ND. <em>Zonula Occludens</em> Toxin Increases the Permeability of Molecular Weight Markers and Chemotherapeutic Agents Across the Bovine Brain Microvessel Endothelial Cells. Journal of Pharmaceutical Sciences. 2003;92(2):414–23.

48. Burkhart A, Helgudóttir SS, Mahamed YA, Fruergaard MB, Holm-Jacobsen JN, Haraldsdóttir H, et al. Activation of glial cells induces proinflammatory properties in brain capillary endothelial cells in vitro. Scientific Reports. 2024;14(1):26580.

49. Voirin A-C, Perek N, Roche F. Inflammatory stress induced by a combination of cytokines (IL-6, IL-17, TNF-α) leads to a loss of integrity on bEnd.3 endothelial cells in vitro BBB model. Brain Research. 2020;1730:146647.

50. Vance ML, Nagy D, Brunner E, Morkotinis V, Black JL, Refai LH, et al. Endothelial-to-mesenchymal transition in the central nervous system: A potential therapeutic target to combat age-related vascular fragility. The Journal of Pharmacology and Experimental Therapeutics. 2025;392(11).

51. Dejana E, Hirschi KK, Simons M. The molecular basis of endothelial cell plasticity. Nature Communications. 2017;8(1):14361.

52. Hartrampf S, Dudakov JA, Johnson LK, Smith OM, Tsai J, Singer NV, et al. The central nervous system is a target of acute graft versus host disease in mice. Blood. 2013;121(10):1906–10.

53. Pinals RL, Islam MR, King O, Choi A, Kang E, Nakano M, et al. Inflammatory reprogramming of human brain endothelial cells compromises blood–brain barrier integrity in Alzheimer’s disease. bioRxiv. 2025:2025.09.26.678918.

54. Sun Z, Zhao H, Fang D, Davis CT, Shi DS, Lei K, et al. Neuroinflammatory disease disrupts the blood-CNS barrier via crosstalk between proinflammatory and endothelial-to-mesenchymal-transition signaling. Neuron. 2022;110(19):3106–20.e7.

55. Luo Y, Yang H, Wan Y, Yang S, Wu J, Chen S, et al. Endothelial ETS1 inhibition exacerbate blood–brain barrier dysfunction in multiple sclerosis through inducing endothelial-to-mesenchymal transition. Cell Death & Disease. 2022;13(5):462.

56. Derada Troletti C, Fontijn RD, Gowing E, Charabati M, van Het Hof B, Didouh I, et al. Inflammation-induced endothelial to mesenchymal transition promotes brain endothelial cell dysfunction and occurs during multiple sclerosis pathophysiology. Cell Death & Disease. 2019;10(2):45.

57. van der Velden WJ, Herbers AH, Feuth T, Schaap NP, Donnelly JP, Blijlevens NM. Intestinal damage determines the inflammatory response and early complications in patients receiving conditioning for a stem cell transplantation. PLoS One. 2010;5(12):e15156.

58. Jansen SA, Nieuwenhuis EES, Hanash AM, Lindemans CA. Challenges and opportunities targeting mechanisms of epithelial injury and recovery in acute intestinal graft-versus-host disease. Mucosal Immunology. 2022;15(4):605–19.

59. Janelsins MC, Lei L, Netherby-Winslow C, Kleckner AS, Kerns S, Gilmore N, et al. Relationships between cytokines and cognitive function from pre- to post-chemotherapy in patients with breast cancer. Journal of Neuroimmunology. 2022;362.

60. Yap NY, Toh YL, Tan CJ, Acharya MM, Chan A. Relationship between cytokines and brain-derived neurotrophic factor (BDNF) in trajectories of cancer-related cognitive impairment. Cytokine. 2021;144:155556.

61. Cheung YT, Ng T, Shwe M, Ho HK, Foo KM, Cham MT, et al. Association of proinflammatory cytokines and chemotherapy-associated cognitive impairment in breast cancer patients: a multi-centered, prospective, cohort study†. Annals of Oncology. 2015;26(7):1446–51.

62. Min CK, Lee WY, Min DJ, Lee DG, Kim YJ, Park YH, et al. The kinetics of circulating cytokines including IL-6, TNF-α, IL-8 and IL-10 following allogeneic hematopoietic stem cell transplantation. Bone Marrow Transplantation. 2001;28(10):935–40.

63. Wang XS, Shi Q, Williams LA, Cleeland CS, Mobley GM, Reuben JM, et al. Serum interleukin-6 predicts the development of multiple symptoms at nadir of allogeneic hematopoietic stem cell transplantation. Cancer. 2008;113(8):2102–9.

64. Farias MG, de Mello Vicente B, Habigzang M, Hirakata VN, de Oliveira da Silva P, Paz AA, Daudt LE. High plasma IL-6 levels following haploidentical allogeneic hematopoietic stem cell transplantation post-transplant cyclophosphamide as predictor of early death and worse outcome. Transplant Immunology. 2022;71:101543.

65. Zhang P, Fleming P, Andoniou CE, Waltner OG, Bhise SS, Martins JP, et al. IL-6–mediated endothelial injury impairs antiviral humoral immunity after bone marrow transplantation. The Journal of Clinical Investigation. 2024;134(7).

66. Arpaia N, Campbell C, Fan X, Dikiy S, van der Veeken J, deRoos P, et al. Metabolites produced by commensal bacteria promote peripheral regulatory T-cell generation. Nature. 2013;504(7480):451–5.

67. Meyer F, Seibert FS, Nienen M, Welzel M, Beisser D, Bauer F, et al. Propionate supplementation promotes the expansion of peripheral regulatory T-Cells in patients with end-stage renal disease. J Nephrol. 2020;33(4):817–27.

68. Nastasi C, Fredholm S, Willerslev-Olsen A, Hansen M, Bonefeld CM, Geisler C, et al. Butyrate and propionate inhibit antigen-specific CD8(+) T cell activation by suppressing IL-12 production by antigen-presenting cells. Sci Rep. 2017;7(1):14516.

69. Masetti R, Leardini D, Muratore E, Fabbrini M, D’Amico F, Zama D, et al. Gut microbiota diversity before allogeneic hematopoietic stem cell transplantation as a predictor of mortality in children. Blood. 2023;142(16):1387–98.

70. Biagi E, Zama D, Nastasi C, Consolandi C, Fiori J, Rampelli S, et al. Gut microbiota trajectory in pediatric patients undergoing hematopoietic SCT. Bone Marrow Transplantation. 2015;50(7):992–8.

71. Romick-Rosendale LE, Haslam DB, Lane A, Denson L, Lake K, Wilkey A, et al. Antibiotic Exposure and Reduced Short Chain Fatty Acid Production after Hematopoietic Stem Cell Transplant. Biology of Blood and Marrow Transplantation. 2018;24(12):2418–24.

72. Cross C, Davies M, Bateman E, Crame E, Joyce P, Wignall A, et al. Fibre-rich diet attenuates chemotherapy-related neuroinflammation in mice. Brain, Behavior, and Immunity. 2024;115:13–25.

73. Mann ER, Lam YK, Uhlig HH. Short-chain fatty acids: linking diet, the microbiome and immunity. Nature Reviews Immunology. 2024;24(8):577–95.

74. Peterson CT, Perez Santiago J, Iablokov SN, Chopra D, Rodionov DA, Peterson SN. Short-Chain Fatty Acids Modulate Healthy Gut Microbiota Composition and Functional Potential. Current Microbiology. 2022;79(5):128.

75. De Filippis F, Pellegrini N, Vannini L, Jeffery IB, La Storia A, Laghi L, et al. High-level adherence to a Mediterranean diet beneficially impacts the gut microbiota and associated metabolome. Gut. 2016;65(11):1812–21.

76. Lippmann ES, Azarin SM, Palecek SP, Shusta EV. Commentary on human pluripotent stem cell-based blood–brain barrier models. Fluids and Barriers of the CNS. 2020;17(1):64.

77. Dei Cas M, Paroni R, Saccardo A, Casagni E, Arnoldi S, Gambaro V, et al. A straightforward LC-MS/MS analysis to study serum profile of short and medium chain fatty acids. J Chromatogr B Analyt Technol Biomed Life Sci. 2020;1154:121982.

78. Hollmann EK, Bailey AK, Potharazu AV, Neely MD, Bowman AB, Lippmann ES. Accelerated differentiation of human induced pluripotent stem cells to blood-brain barrier endothelial cells. Fluids Barriers CNS. 2017;14(1):9.

79. Ebert LM, Tan LY, Johan MZ, Min KKM, Cockshell MP, Parham KA, et al. A non-canonical role for desmoglein-2 in endothelial cells: implications for neoangiogenesis. Angiogenesis. 2016;19(4):463–86.

