## Supplementary Fig.1 for "Gut-immune signaling drives blood-brain barrier damage in pediatric allogeneic stem cell transplant"

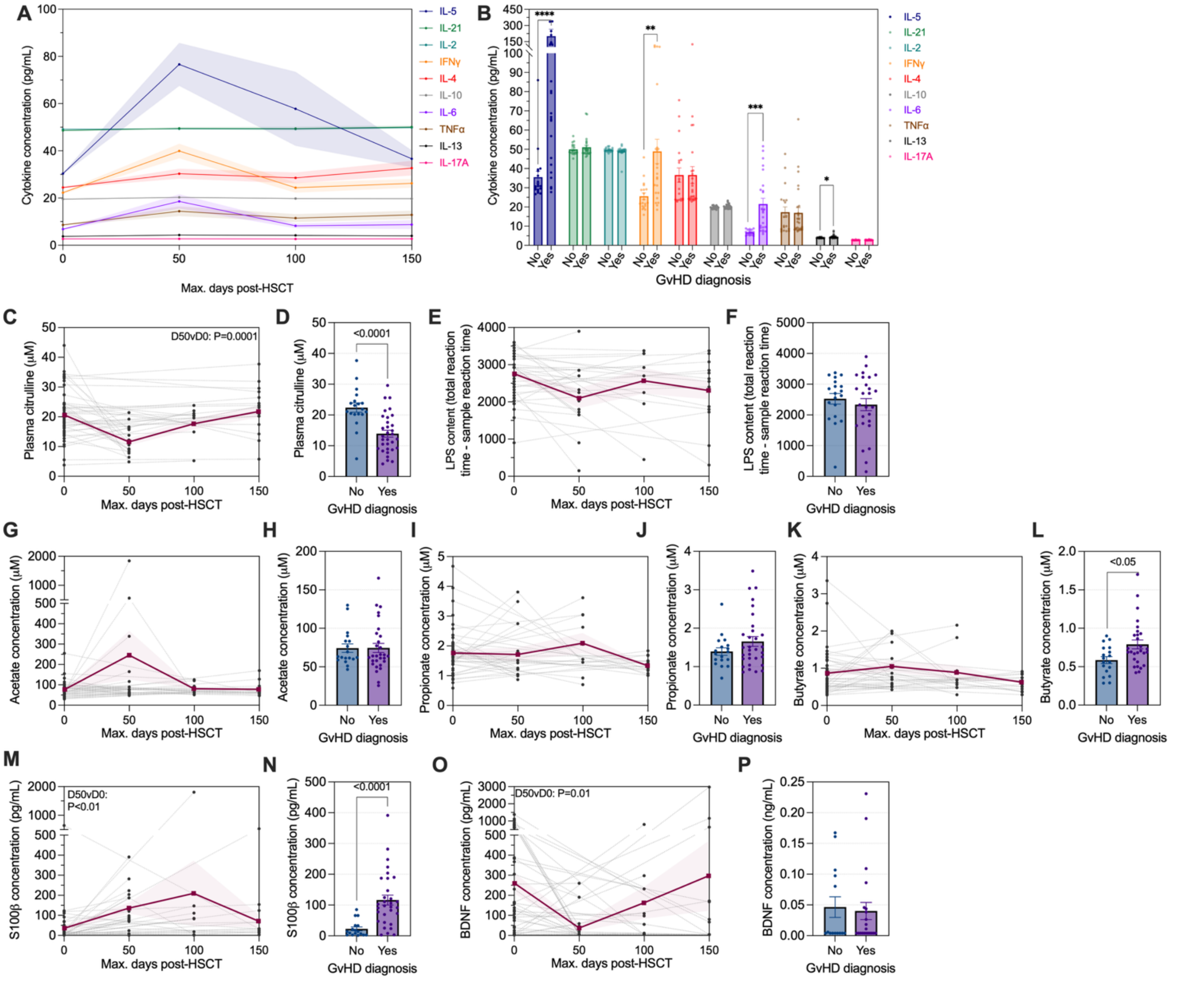

**Supplementary Fig.1 | Longitudinal biomarker and GvHD sub-group analysis**

Longitudinal analysis of biomarkers binned into 50-day intervals (D0: N=47; D1-50: N=17; D51-100: N=10; D101-150: N=18), and post-transplant characterization of biomarkers sub-grouped into GvHD diagnosis status (No: N=19; Yes: N=32). For longitudinal analysis a mixed-effects analysis was performed relative to baseline and for histogram data a Mann-Whitney test was performed for statistical analysis.

(A and B) Cytokines

(C and D) Citrulline

(E and F) LPS

(G and H) Acetate

(I and J) Propionate

(K and L) Butyrate

(M and N) S100β

(O and P) BDNF

**p<0.01, ***p<0.001. Histogram data and longitudinal data for cytokines represent mean ±SEM. Longitudinal data for all other displayed as individual patient trajectories represented by grey dotted lines and the overall mean depicted in bold, maroon line overlayed.
