## Supplementary Fig.2 for "Gut-immune signaling drives blood-brain barrier damage in pediatric allogeneic stem cell transplant"

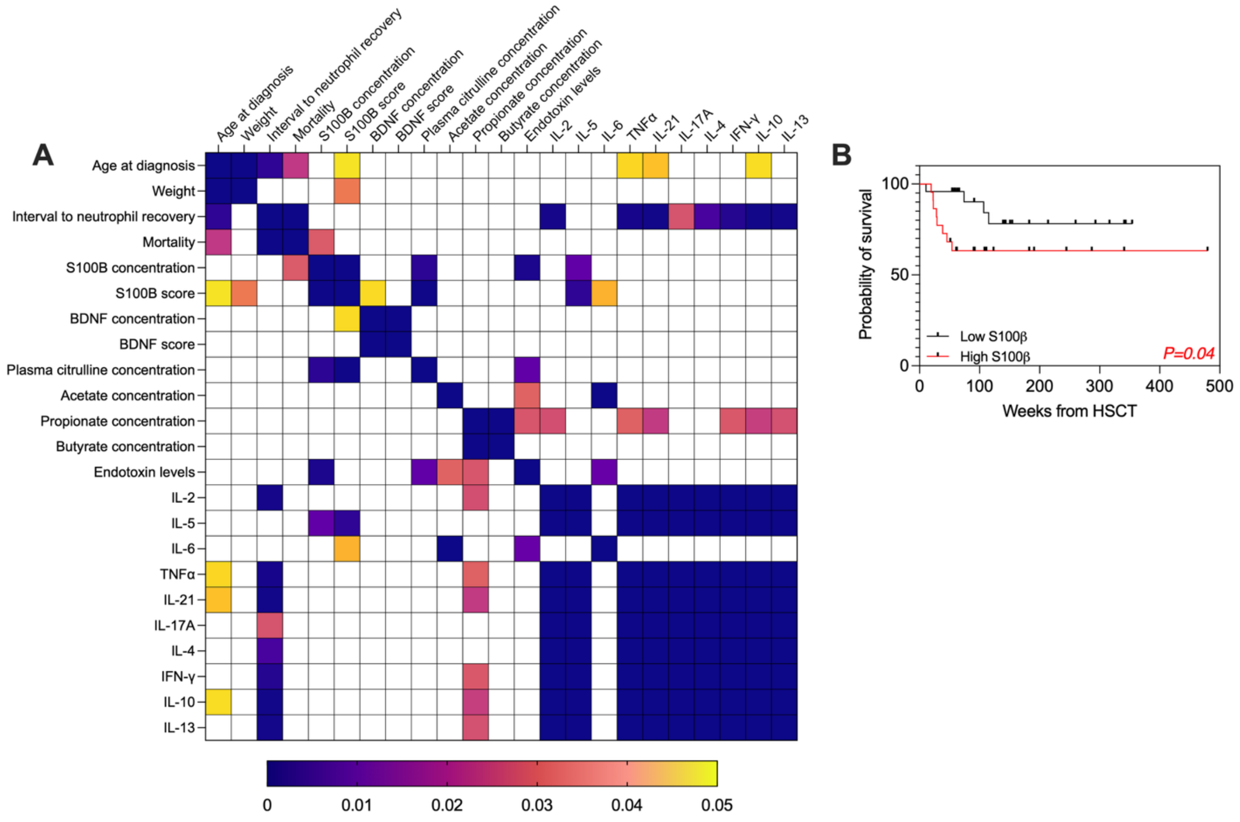


**Supplementary Fig.2 | Significant correlation matrix and S100β survival analysis**

(A) Correlation matrix heatmap indicating only the significant relationships, P-values determined via Pearson's correlation and signified by grid square color.

(B) Plasma samples post-HSCT categorized as high or low S100β if higher or lower than the cohort median. Comparison of probability of survival between high and low groups was performed via Gehan-Breslow-Wilcoxon test.
