## Supplementary Fig.3 for "Gut-immune signaling drives blood-brain barrier damage in pediatric allogeneic stem cell transplant"

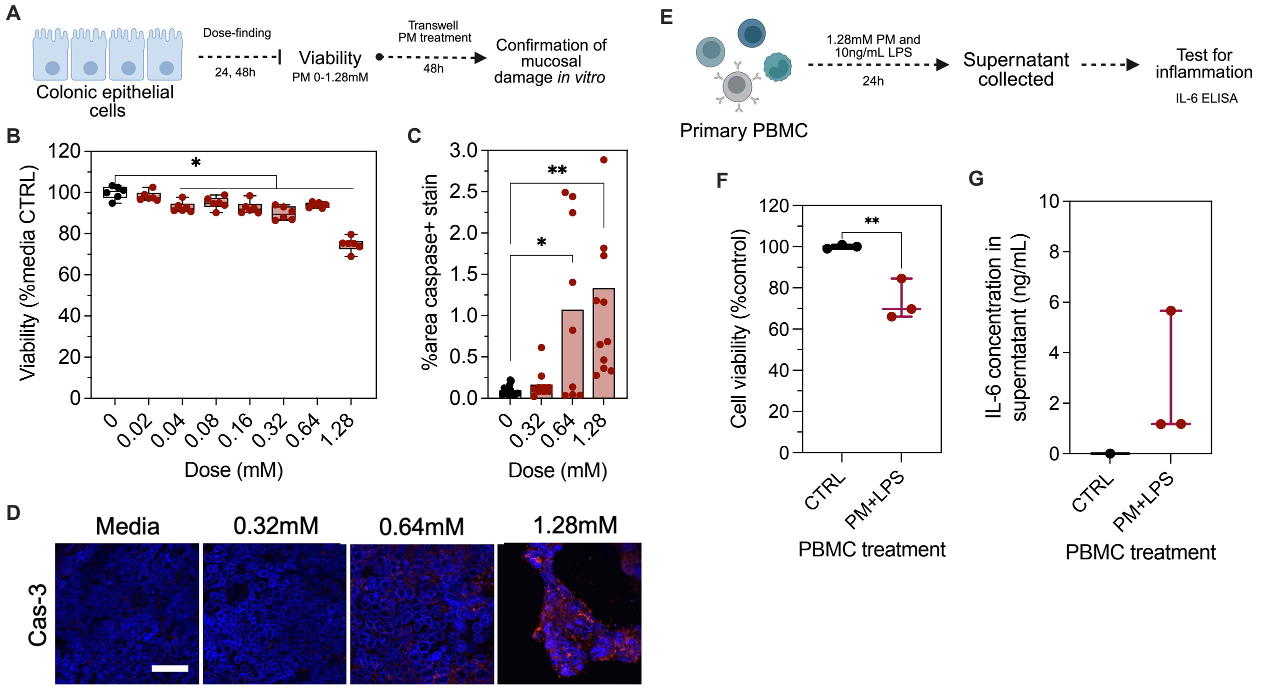


**Supplementary Fig.3 | PM dose-finding in T84 epithelial cells and PBMC stimulation modelling cyclophosphamide-induced mucositis immune activation.**

(A) Experimental design of PM dose-finding with T84 colonic epithelial cells.

(B) Results of XTT-assay of T84 viability (% media only control) at 48-hours (one-way ANOVA; N=6 wells of a 96-well plate per condition).

(C) Caspase-3 positive area staining after 48-hour exposure to PM or media only control (one-way ANOVA; N=4 transwells per condition).

(D) Representative images of T84 caspase-3 staining (scale bar indicated 30um).

(E) Experimental design of PBMC stimulation with 1.28mM PM and 10ng/mL LPS (or PBS control) for 24h.

(F) PBMC viability post-stimulation relative to baseline (control) viability (unpaired t-test; N=3).

(G) IL-6 concentrations in PBMC stimulation (PBS CTRL: N=3; PM+LPS: N=1).

*p<0.05, **p<0.01. Box and whisker plots represent median and IQRs (B), or median (F and G).
