## Supplementary Fig.4 for "Gut-immune signaling drives blood-brain barrier damage in pediatric allogeneic stem cell transplant"

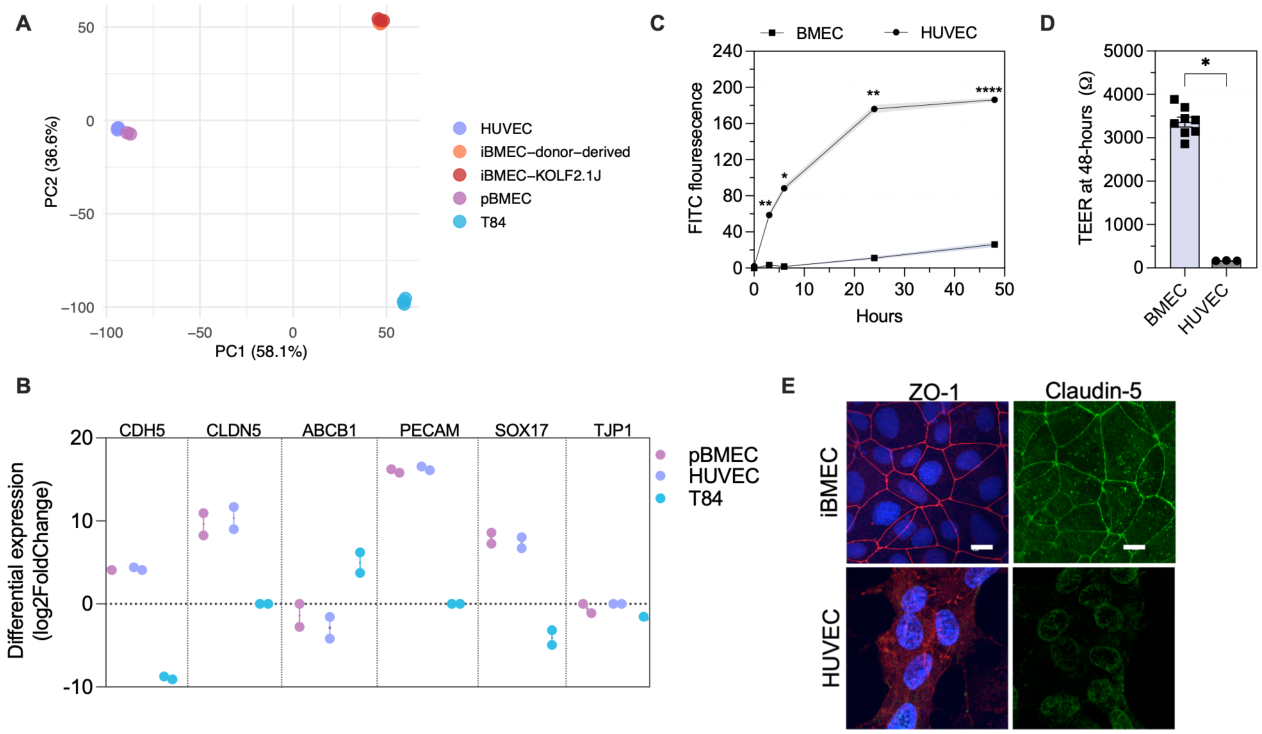


**Supplementary Fig.4 | Evaluation of iPSC-derived BMECs**

(A-D) Bulk RNA sequencing comparison of iPSC-derived BMECs (iBMECs) derived from two iPSC sources (donor-derived or KOLF2.1J) to primary BMECs (pBMECs), immortalized human umbilical vein endothelial cells (HUVECs), or T84 colonic epithelial cells (N=2 for pBMECs, N=3 for all other cell types; ~33.2 million paired-end reads per sample).

(A) Principal component analysis.

(B) Differential expression of key endothelial markers by comparator cells relative to the two sources of iBMECs, fold-change of 0 indicative of non-significance.

(C) Comparison of 4kDA FITC translocation across a monolayer of iBMECs (donor-derived) vs HUVECs, cultured on transwells (mixed-effects analysis; N=3 transwells).

(D) Trans-endothelial electrical resistance (TEER) values measured 48-hours after transwell subculture (unpaired t-test; HUVEC N=3, BMEC N=8),

(E) Immunofluorescent visualization of tight junction (ZO-1 and claudin-5) formation (scale bar indicates 15μm).

*p<0.05, **p<0.01, ****p<0.0001. (C and D) represent mean ±SEM.
