## Supplementary Fig.5 for "Gut-immune signaling drives blood-brain barrier damage in pediatric allogeneic stem cell transplant"

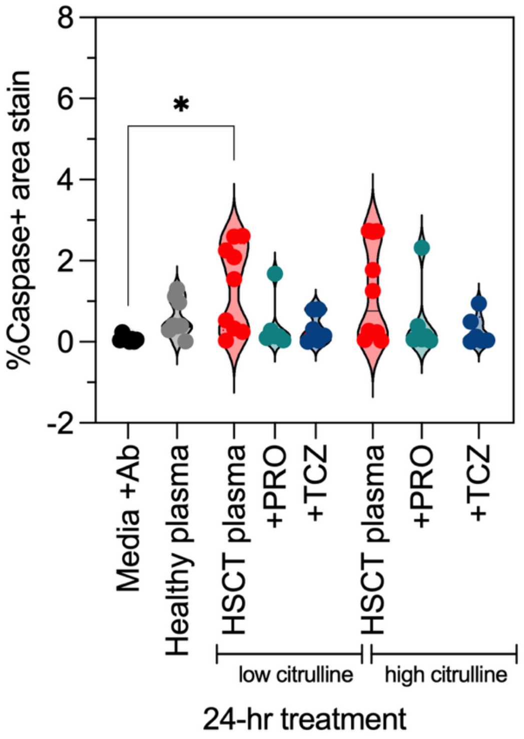


**Supplementary Fig.5 | Allo-HSCT plasma-induced iBMEC apoptosis stratified by patient citrulline levels**

Caspase-3 positive %area stain 24-hr after exposure to patient plasma (Kruskal-Wallis test; N=2 transwells per condition and a minimum N=6 FOVs for caspase-3 quantification).

*$p<0.05$, violin plot denotes median and min-max values.
